# Functional modulation of PTH1R activation and signalling by RAMP2

**DOI:** 10.1101/2021.12.08.471790

**Authors:** Katarina Nemec, Hannes Schihada, Gunnar Kleinau, Ulrike Zabel, Eugene O. Grushevskyi, Patrick Scheerer, Martin J. Lohse, Isabella Maiellaro

## Abstract

Receptor-activity-modifying proteins (RAMPs) are ubiquitously expressed membrane proteins that associate with different G protein-coupled receptors (GPCRs) including the parathyroid hormone 1 receptor (PTH1R), a class B GPCR and an important modulator of mineral ion homeostasis and bone metabolism. However, it is unknown whether and how RAMP proteins may affect PTH1R function.

Using different optical biosensors to measure the activation of PTH1R and its downstream signalling, we describe here that RAMP2 acts as a specific allosteric modulator of PTH1R, shifting PTH1R to a unique pre-activated state that permits faster activation in a ligand-specific manner. Moreover, RAMP2 modulates PTH1R downstream signalling in an agonist-dependent manner, most notably increasing the PTH-mediated Gi3 signalling sensitivity. Additionally, RAMP2 increases both PTH- and PTHrP-triggered β-arrestin2 recruitment to PTH1R. Employing homology modelling we describe the putative structural molecular basis underlying our functional findings.

These data uncover a critical role of RAMPs in the activation and signalling of a GPCR that may provide a new venue for highly specific modulation of GPCR function and advanced drug design.

## Introduction

G protein–coupled receptors (GPCRs) represent the largest class of membrane-bound proteins and are involved in a multitude of biological processes^1^. They are characterized by a seven-transmembrane helix structure, which undergoes a characteristic rearrangement upon binding of agonists. Agonist binding to its cognate receptor induces conformational changes in the transmembrane helices, which are transmitted to the cytosolic face of the receptors and ultimately result in receptor activation, which represents the key step of signal transduction. The combination of crystallographic and cryo-EM studies and the employment of optical biosensors to study the reorganization of the seven transmembrane domains has allowed a detailed understanding of the general mechanisms of GPCR receptor activation^2-5^.

Earlier structural studies suggest that GPCRs undergo similar conformational changes upon activation including, most prominently, an outward movement of the transmembrane helix 6 at the cytosolic face, thereby creating a pocket to which the G protein α-subunit can couple^5^. More recent studies, however, have revealed that the exact type of changes may depend on the receptor class and the specific receptor^6,7,8^. Class-and receptor-specific differences may also exist in the interaction of receptors not only with downstream G proteins and β-arrestins, but also with accessory and modulatory proteins^9^.

Studies of the kinetic steps that govern the structural rearrangements which underlie receptor activation^10^ showed that its speed may depend on the receptor class and the specific receptor. When exposed to saturating agonist concentrations, most class A GPCRs switch into the active state within tens of milliseconds, while early steps of class C receptor activation have been observed to occur within 1-2 ms, and activation of class B receptors may take much longer^11-15^. Little is known whether the activation kinetics of GPCRs can be modulated by their cellular context and whether proteins other than the receptors themselves might play a role in shaping signalling kinetics and specificity.

Here, we study the parathyroid hormone 1 receptor (PTH1R), a prototypical member of class B GPCRs, which are characterized by a large N-terminal domain that binds a major part of their cognate peptide agonists^16, 17^. Compared to class A GPCRs, PTH1R activation is relatively slow and occurs in a two-step process: the initial N-terminal binding step has a time constant of ≈140 ms and is followed by an interaction of the ligand with the transmembrane core, which changes into its active conformation with a time constant of ≈1 s^11,14^. Pleiotropic in its downstream coupling, PTH1R signals primarily via G_s_, but can also couple to G_q_^18^, G_12/13_ ^19^ and G_i_ ^20^, and interacts with and signals via β-arrestins ^21,22^. The two endogenous agonists, parathyroid hormone (PTH) and parathyroid hormone-related peptide (PTHrP), trigger PTH1R activation with similar kinetics and specificity for the various intracellular pathways^23-25^. However, PTH can induce prolonged signalling from intracellular sites, while PTHrP signals exclusively from the cell surface^26^. PTH1R has been reported to interact with modulatory proteins of the receptor-activity-modifying protein (RAMP) family^27-29^. RAMPs constitute a family of single transmembrane helix proteins with three members: RAMP1, RAMP2, and RAMP3.

It is controversial whether PTH1R interacts only or preferentially with RAMP2^27^ or all three RAMPs^28,29^. In RAMP2 knock-out mice, PTH1R function is deregulated, and placental dysfunction is observed^30^, suggesting a major physiological role of the PTH1R/RAMP2 interaction. However, the molecular mechanisms how RAMPs may modulate the activation dynamics of PTH1R and their signalling properties remain to be elucidated.

To address these questions, we develop and employ new biosensors for PTH1R activation and investigate a set of downstream signalling pathways to assess the effects of RAMPs on the activation dynamics and signalling properties of PTH1R in response to its two endogenous ligands, PTH and PTHrP. We observe that RAMP2 specifically interacts with PTH1R and modulates its activation kinetics as well as signalling dynamics in an agonist-dependent manner.

## Results

### Analysis of PTH1R/RAMP interactions at the cell surface

First, we investigated the interactions of PTH1R with the three RAMPs at the surface of intact cells. To do so, we performed acceptor photobleaching experiments to quantify FRET efficiencies between PTH1R, labelled with a C-terminal mCitrine^31^ (mC), and the three different RAMPs, labelled at their intracellular C-terminus with mTurquoise2^32^ (mT2) (**Fig. 1*A***). FRET efficiencies were quantified by measuring the recovery of donor emission after photobleaching of the acceptor (**Fig. 1*B***) in human embryonic kidney (HEK293) cells transiently co-expressing comparable levels of tagged PTH1R in combination with tagged RAMP1, -2 or -3 (*SI Appendix*, Fig. S1A).

**Fig. 1.**
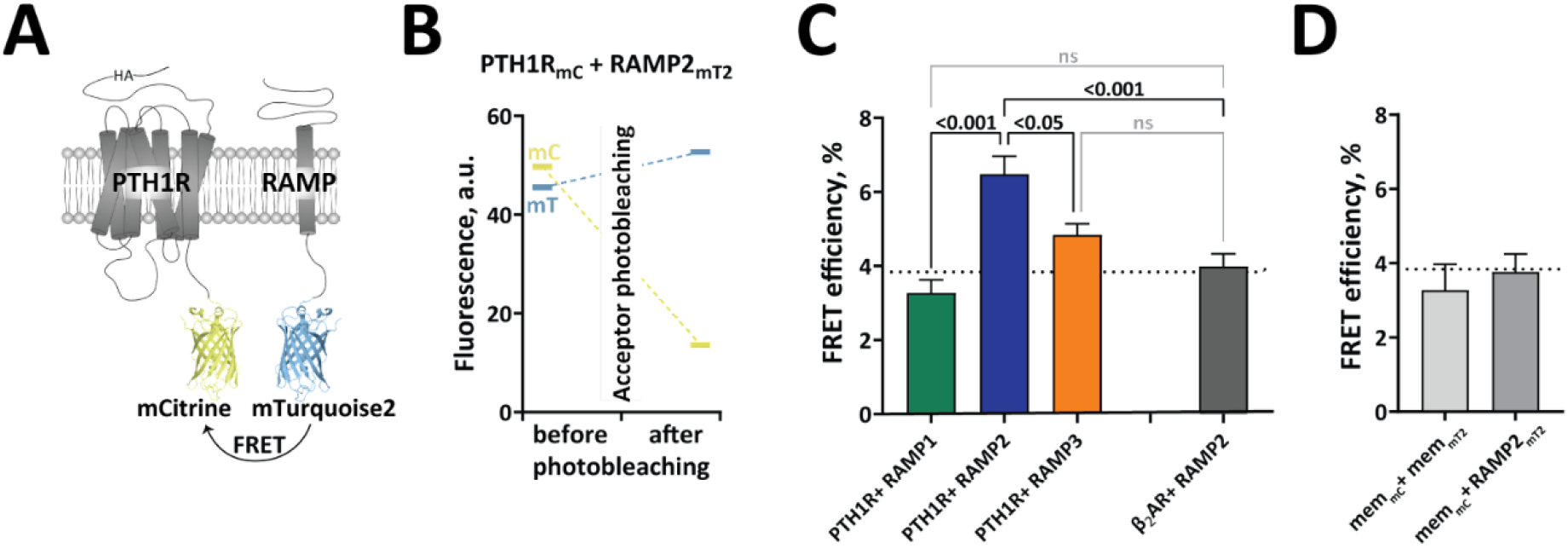
Intermolecular FRET reveals RAMP2 as an interaction partner of PTH1R. (A) Schematic representation of the constructs for FRET acceptor photobleaching experiments between PTH1R and RAMP2. Photobleaching experiments were conducted in HEK293 cells, transiently co-transfected with a combination of donor- and acceptor-tagged constructs. The acceptor fluorophore (mCitrine, mC) was fused to the C-terminal of the PTH1R (PTH1R_mC_), to the control β_2_-adrenergic receptor (β_2_AR_mC_) or targeted to the plasma membrane via a -CAAX sequence (mem_mC_). The donor fluorophore (mTurquoise2, mT2) was fused to the C-terminal of RAMPs (RAMP1/2/3_mT2_) or targeted to the plasma membrane via a -CAAX sequence (mem_mT2_). (B) Representative experiment showing the efficiency of photobleaching in cells expressing PTH1R_mC_ and RAMP2_mT2_. Fluorescence emission of both donor (mC) and acceptor (mT2) were recorded before and after acceptor photobleaching. (C, D) FRET efficiencies from photobleaching experiments recorded with a confocal microscope. The data are expressed as % of donor emission increase after photobleaching for each experimental group. The dotted line indicates the average FRET efficiency of negative control groups (grey bars). The data are derived from at least 3 independent experiments and following numbers of cells: PTH1R_mC_ + RAMP1_mT2_ (n = 46), PTH1R_mC_ + RAMP2_mT2_ (n = 70), PTH1R_mC_ + RAMP3_mT2_ (n = 71), β_2_AR_mC_ + RAMP2_mT2_ (n = 51), mem_mC_ + RAMP2_mT2_ (n = 37) and mem_mC_ + mem_mT2_ (n = 9); bars represent means ± SEM. Significance between the groups was assessed by Brown-Forsythe ANOVA, followed by Dunnett’s multiple comparisons test, ns: p > 0.05.

The FRET efficiency was significantly higher for cells expressing PTH1R_mC_ with RAMP2_mT2_ than for combinations with either RAMP1_mT2_ or RAMP3_mT2_ (**Fig. 1*C***). In fact, the FRET efficiencies for the latter two were not significantly different from background FRET (p>0.05; dotted line in **Fig. 1*C* and *D***), which is determined by either non-specific FRET between two membrane tags or between a membrane tag and RAMP2_mT2_ (**Fig. 1*D***), or by FRET between RAMP2_mT2_ and the β_2_-adrenergic receptor (β_2_AR_mC_), a GPCR shown not to interact with RAMP2^29,33^. These data indicate that PTH1R forms complexes with RAMP2 at the cell surface, but very little or none with RAMP3 or RAMP1.

### RAMP2 expression modulates PTH1R basal and PTH-bound conformations

We then aimed to investigate whether RAMP2 regulates PTH1R activation dynamics. Based on previously reported PTH1R biosensors with donor and acceptor fluorophores fused to conformationally sensitive sites^11,34^, we generated an improved conformational biosensor, PTH1R_FRET_. Preserving insertion sites in the third intracellular loop and at the C-terminus, we exchanged the fluorophores with brighter and more photostable fluorophores, namely mT2 and mC (**Fig. 2*A***).

**Fig. 2.**
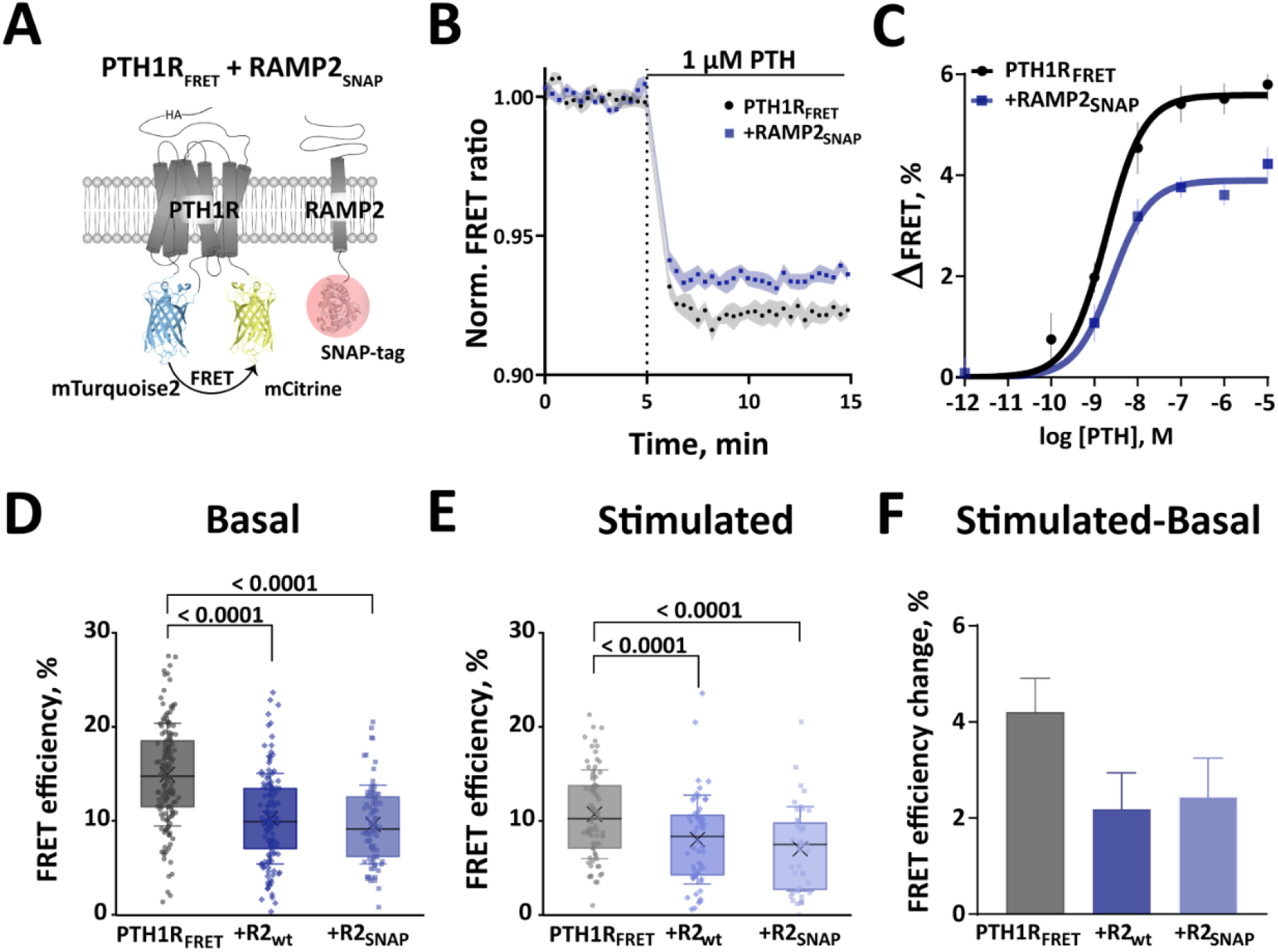
RAMP2 modulates PTH1R basal and PTH-induced conformations. (A) Schematic representation of intramolecular PTH1R_FRET_ biosensor. To control for RAMP2 expression, its C-terminally fused SNAP-tag was labelled with 1 µM SNAP-Cell SiR-647 (red circle). (B) Average time course of PTH-induced FRET changes recorded in a plate reader from HEK293 cells transiently expressing PTH1R_FRET_ (black) alone or together with RAMP2_SNAP_ (blue). The data are an average of five independent experiments normalized to the initial FRET value (set to 1). (C) Concentration-response curves obtained from traces as in (B). ΔFRET values are expressed as percent change from the initial FRET value. Curve fitting gave pEC_50_ values (means ± SEM) of: PTH1R_FRET_ = 8.73 ± 0.12 and PTH1R_FRET_ + RAMP2_SNAP_ = 8.59 ± 0.17. (D, E) FRET efficiencies from photobleaching experiments recorded with a confocal microscope. The data are expressed as % of donor emission increase after photobleaching for each experimental group. FRET efficiencies of basal (D) and 100 µM PTH-stimulated (E) HEK293 cells transiently expressing PTH1R_FRET_ (black) alone or together with RAMP2_wt_ (dark blue) or RAMP2_SNAP_ (light blue). The data are from the following numbers of cells obtained in six (basal) and three (stimulated) independent experiments. Basal: PTH1R_FRET_ (n = 153), PTH1R_FRET_ + RAMP2_wt_ (n = 130), PTH1R_FRET_ + RAMP2_SNAP_ (n = 82). Stimulated PTH1R_FRET_ (n = 73), PTH1R_FRET_ + RAMP2_wt_ (n = 56), PTH1R_FRET_ + RAMP2_SNAP_ (n = 44). Data show values from individual cells; boxes represent the first and third interquartile range, and whiskers indicate SD. Significance between the groups was tested with one-way ANOVA followed by Dunnett’s multiple comparisons test, ns: p > 0.05. (F) FRET efficiency changes calculated from (D) and (E), represented as percent change ± SEM.

To measure PTH1R activation dynamics, we co-transfected HEK293 cells with PTH1R_FRET_ biosensor with and without RAMP2_SNAP_. To estimate the expression level of RAMP2 in these experiments, we tagged the C-terminus of RAMP2 with a SNAP-tag which was labelled with the permeable fluorescent dye SNAP-Cell SiR-647^35^. This allowed us to determine that PTH1R expression was not affected by up to 2 µg of RAMP2_SNAP_ cDNA (corresponding to a 1:2 PTH1R:RAMP2 transfection ratio), and a 1:1 transfection ratio (PTH1R:RAMP2) was used for all subsequent experiments (*SI Appendix*, Fig. S2 A and B). Further control experiments showed that neither cell surface expression of PTH1R_FRET_ (measured with ELISA via detection of a HA-tag in the PTH1R_FRET_; *SI Appendix*, Fig. S2C), nor the total expression of PTH1R_FRET_ (measured by direct excitation of mC in the PTH1R_FRET_; *SI Appendi*x, Fig. S2D) were affected by the expression of RAMP2, either in its wildtype or in its SNAP-tagged form.

We then measured the changes in FRET of PTH1R_FRET_ evoked by different concentrations of PTH, initially by manual addition in a microtiter plate format. As in similar GPCR biosensors, agonists evoked a decrease of FRET, presumably induced by a movement of the third intracellular loop away from the C terminus, which is thought to cause an increased distance between the two fluorophores in the biosensor^11,15-17^. Figure 2B shows the time courses of the PTH-induced decrease in the FRET ratio in control (black) and cells, co-expressing RAMP2 (blue). The amplitude of this decrease was smaller in RAMP2_SNAP_ expressing cells than in control cells at all concentrations of PTH (**Fig. 2 *B,C***), while the potencies of PTH were not different between the two conditions (**Fig. 2*C***); these effects were the same whether SNAP-tagged or wildtype RAMP2 was used (*SI Appendix*, Fig. S2E).

To assess whether the decrease in the amplitude of the PTH-induced FRET signal by RAMP2 might be caused by a change in the initial conformation and, hence, basal FRET of the biosensor, we performed photobleaching experiments of PTH1R_FRET_ in the absence or presence of RAMP2_SNAP_. We observed that under basal conditions the FRET efficiency was significantly higher in the absence than in the presence of RAMP2, with no difference between C-terminally labelled RAMP2_SNAP_ or wildtype RAMP2 (**Fig. 2*D***).

After 5 min of stimulation with a high concentration (100 µM) of PTH, a similar pattern was observed, i.e. the FRET efficiency was higher in the absence than in the presence of RAMP2 (**Fig. 2*E*)**. Again, there was no difference between wildtype RAMP2 and RAMP2_SNAP_, indicating that the two could be used interchangeably, and that a SNAP-tag on the C-terminus did not affect the effect of RAMP2 on PTH1R. Interestingly, the RAMP2-induced decrease in FRET efficiency was smaller in the PTH-activated state than under basal conditions (**Fig. 2*E* vs. 2*D***), and similarly the PTH-induced decrease in FRET was smaller in the presence of RAMP2 than in its absence (**Fig. 2*F***).

Control experiments showed that PTH1R_FRET_ biosensor expression was comparable across all tested groups (*SI Appendix*, Fig. S3A), as was the amount of bleaching in each group (*SI Appendix*, Fig. S3B). Additionally, a hyperbolic increase of FRET efficiencies at increasing concentrations of acceptor as determined by prebleached emissions demonstrated that FRET was specific^15^ (*SI Appendix*, Fig. S3C).

Taken together, these data indicate that RAMP2 modulates the conformation of the PTH1R_FRET_ biosensor: it decreases FRET in the basal state and, less so, in the PTH-activated state, and it decreases the PTH-induced FRET signal. A possible explanation for these findings is that RAMP2 induces a kind of pre-activation of PTH1R_FRET_, which is characterized by decreased basal FRET.

### RAMP2 modulates the activation speed and the amplitude of the PTH1R_FRET_ biosensor

To assess whether the interaction of RAMP2 with PTH1R might modulate the activation kinetics we performed experiments with a rapid superfusion system^11^, using a stable cell line expressing PTH1R_FRET_ (**Fig. 2*A***). Transient co-expression and visualization of RAMP2_SNAP_ were performed as described above.

Stimulation with a saturating concentration of PTH (10 µM), in the presence and absence of RAMP2_SNAP,_ resulted in a rapid decrease of the FRET ratio (**Fig. 3*A* and *B***), characterized by antiparallel changes of donor and acceptor emission channels (*SI Appendix*, Fig. S4). The amplitude of the FRET change, similar to microtiter plate and photobleaching experiments (**Fig. 2**), was about two-fold higher in the absence than in the presence of RAMP2_SNAP_ (**Fig. 3*D***).

**Fig. 3.**
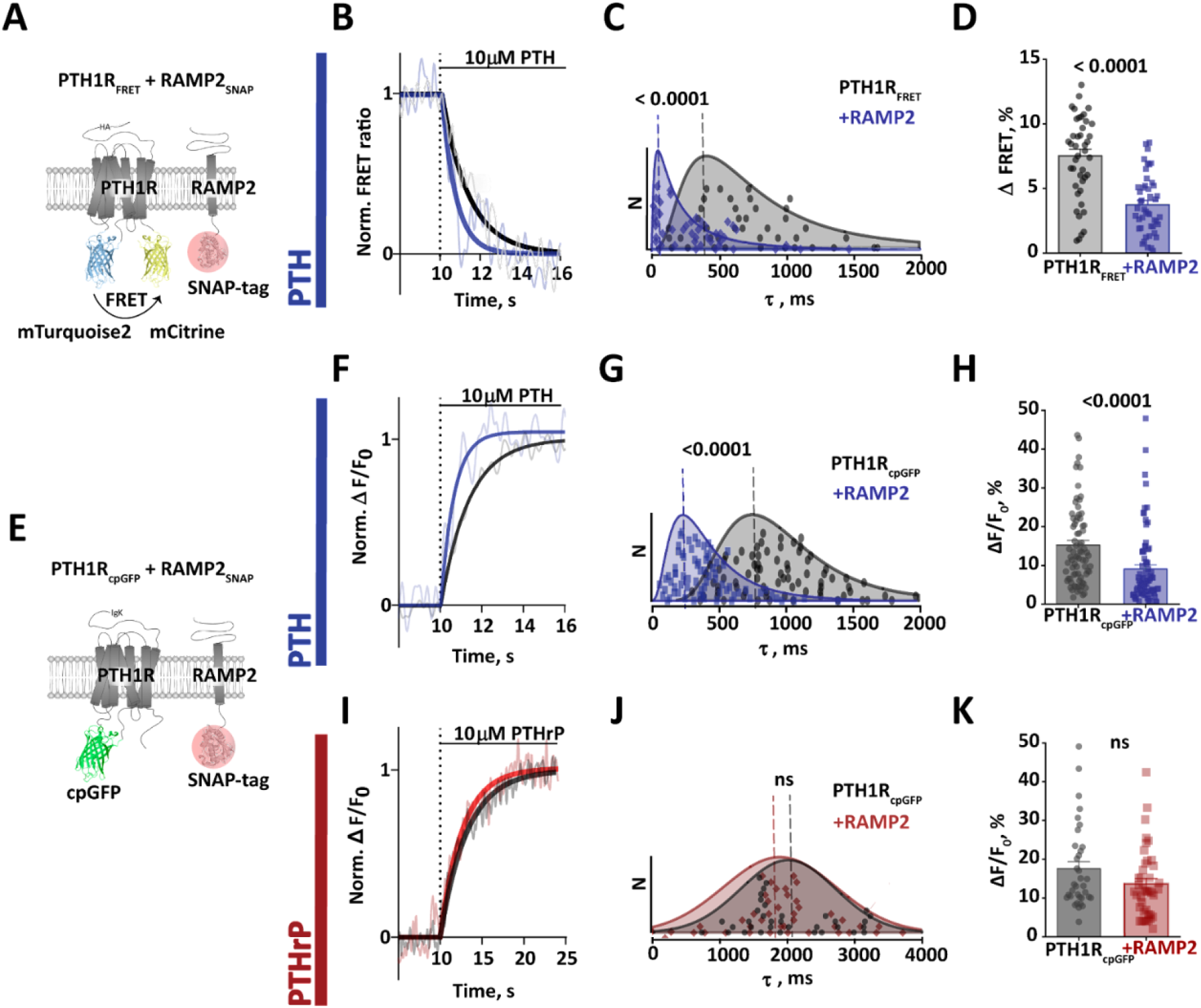
Modulatory effects of RAMP2_SNAP_ co-expression on PTH1R_FRET_ and PTH1R_cpGFP_ biosensor activation. (A) Schematic representation of intramolecular PTH1R_FRET_ biosensor. To control for RAMP2 expression, its C-terminally fused SNAP-tag was labelled with 1 µM SNAP-Cell SiR-647 (red circle). (B) Representative ratio traces of PTH-mediated FRET changes in single HEK293 cells stably expressing PTH1R_FRET_ and in the presence oftransiently co-expressed RAMP2_SNAP_, recorded in a microscopic FRET setup. To analyse only cells that expressed both, cells were labelled with 1 µM SNAP-Cell SiR-647 and regions of interest were selected where PTH1R_FRET_ and RAMP2_SNAP_ were co-expressed. Horizontal lines indicate application of 10 μM PTH with a rapid superfusion system. Traces were normalized to the baseline (set to 1) and plateau after stimulation (set to 0). Shown are FRET ratio traces raw (grey and light blue) and Fourier-lowpassed (black, dark blue). Traces are representative of n = 41 cells (PTH1R_FRET_) and n = 46 cells (+RAMP2_SNAP_), acquired in five independent experiments. (C) Time constants *τ* of PTH-induced FRET changes derived from traces as in panel B, calculated from mono-exponential curve fitting. The data were fitted with a lognormal distribution and the dashed line indicates mode, global maximum of the distribution for: PTH1R_FRET_ = 410 ms; +RAMP2_SNAP_ = 50 ms. Median value and 95% confidence interval (CI) were: PTH1R_FRET_ = 710 ms [516, 946 CI], n = 41 cells; RAMP2_SNAP_ = 330 ms [144, 416 CI], n = 46 cells. A Mann-Whitney test was used to assess a significant difference between the groups (p < 0.001). (D) Amplitude of FRET changes induced by PTH obtained from traces as in panels B. Bars represent means ± SEM, % of the FRET amplitudes from all examined cells: PTH1R_FRET_ = 7.5 ± 0.5 %, PTH1R_FRET_+RAMP2_SNAP_ = 3.8 ± 0.3 %. A t-test was used to assess a significant difference between the groups (p < 0.001). (E) Schematic representation of the single colour biosensor to monitor PTH1R activity in single cell experiments. Receptor activation upon agonist stimulation was monitored by recording fluorescence at 516 nm. (F, I) Representative fluorescence traces of PTH (F) and PTHrP (I) mediated changes in ΔF/F_0_ recorded in a microscopic FRET setup in single HEK293 cells transiently expressing PTH1R_cpGFP_ alone or with RAMP2_SNAP_, labelled with 1 µM SNAP-Cell SiR-647. Horizontal lines indicate application of 10 μM PTH or PTHrP with a rapid superfusion system. Data show are in panel B. (G, J) Time constants *τ* of PTH-induced and PTHrP-induced activation derived from traces as in panels F and I. The data were analyzed as described in panel C. Dashed line indicates mode, global maximum of the distribution: PTH1R_cpGFP_ = 760 ms and PTH1R_cpGFP_+RAMP2_SNAP_ = 190 ms. Median value and 95% confidence intervals for PTH: PTH1R_cpGFP_ = 950 ms [817, 1057 CI], n = 78 cells; PTH1R_cpGFP_+RAMP2_SNAP_ = 400 ms [322, 448 CI], n = 75 cells. A Mann-Whitney test was used to assess a significant difference between the groups (p < 0.001). PTHrP: PTH1R_cpGFP_ = 1960 ms [1770, 2660 CI], n = 38; PTH1R_cpGFP_+RAMP2_SNAP_ = 1910 ms [1670, 2100 CI], n = 41. (H, K) Effects of RAMP2 on the amplitude of the ΔF/F_0_ signals induced by PTH (F) and PTHrP (H). Bars represent means ± SEM in % of the ΔF/F_0_ amplitudes from all cells examined: PTH: PTH1R_cpGFP_ = 15.3 ± 1.1 % (n = 78 cells); PTH1R_cpGFP_+RAMP2_SNAP_ = 9.1 ± 1.1 %, (n = 77 cells). PTHrP: PTH1R_cpGFP_ = 17.5 ± 1.8 % (n = 38 cells); PTH1R_cpGFP_+RAMP2_SNAP_ =13.6 ± 1.4 % (n = 41 cells). A t-test was used to assess a significant difference between the groups (p < 0.001).

Activation time constants (*τ*) were calculated by mono-exponential curve fitting. In accordance with earlier data^11^, PTH activated PTH1R_FRET_ with a median time constant of 710 ms (**Fig. 3*C***). However, when RAMP2_SNAP_ was co-expressed, the PTH-induced activation was twice as fast with a median *τ* value of only 330 ms (**Fig. 3*C***). Figure 3*C* shows the distribution of the time constants under the two conditions, which peak at 410 ms and 50 ms, respectively.

Control experiments showed that neither membrane expression of the PTH1R_FRET_ biosensor measured with ELISA via detection of the HA-tag in PTH1R_FRET_ (*SI Appendix*, Fig. S2C), nor the total expression of PTH1R_FRET_ measured as direct emission of biosensor’s acceptor were affected by RAMP2 (*SI Appendix*, Fig. S2D). This excluded the possibility that differences between the control and RAMP2 group were due to different expression levels of the biosensor.

Overall, these results indicate that RAMP2 has two distinct effects on receptor activation as observed with PTH1R_FRET_: it increases the speed several-fold, and it reduces the amplitude (from a lower starting value) by ≈2-fold.

### A single color PTH1R biosensor confirms the modulatory role of RAMP2

To further substantiate our results, we generated a novel PTH1R biosensor based on a single fluorophore, circularly permuted green fluorescent protein (cpGFP). This approach was originally developed to visualize fast calcium dynamics^36^ and neurotransmitter release – processes that occur with sub-second time courses^37^. We generated a PTH1R_cpGFP_ biosensor by inserting a cpGFP module with linkers into the third intracellular loop (**Fig. 3*E***), similar to the donor insertion position in PTH1R_FRET_. In preliminary experiments in microtiter plate format, PTH1R_cpGFP_ transiently expressed in HEK293 cells showed a marked increase in fluorescence in response to agonist activation, which occurred with a similar potency (*SI Appendix*, Fig. S5B) as in wildtype PTH1R^11^ or PTH1R_FRET_ (**Fig 2*C***, *SI Appendix*, Fig. S2E). Again, we monitored co-expression of RAMP2_SNAP_ via a C-terminal SNAP-tag and analyzed only cells that expressed both, PTH1R_cpGFP_ and RAMP2_SNAP_ (**Fig. 3*E***).

We then performed single cell experiments with these cells, applying agonists via a fast perfusion system. 10 µM PTH evoked an increase in fluorescence in control and RAMP2 expressing cells (**Fig. 3*F***). In agreement with results obtained with the PTH1R_FRET_ biosensor, the amplitude of the signal and also the speed of the activation process were affected by RAMP2: in particular, RAMP2 decreased the amplitude (ΔF/F0) and increased the speed of activation by PTH (**Fig. 3*G* and *H***). The time constant (***τ***) of activation was decreased from 950 ms to 390 ms and the peak of the ***τ***-value distribution from 760 ms to 190 ms by the presence of RAMP2.

Interestingly, similar experiments with the second endogenous agonist, PTHrP, revealed that these effects were agonist-specific: using PTHrP (10 µM) as the agonist, no significant differences in the amplitude (ΔF/F0) or the time constant (***τ***) were detected between control and RAMP2 co-expressing cells (**Fig. 3*I-K***). These results suggest that RAMP2 modulation of PTH1R is agonist-specific.

### Ligand-specific effects of RAMP2 on G protein activation by PTH1R

Since RAMP2 appears to change the PTH1R conformation resulting in faster agonist-specific activation, we wondered whether it might also affect downstream signalling by the PTH1R. To test this hypothesis, we used a suite of BRET- and FRET-based biosensors to quantify the effect of PTH1R activation on different signalling pathways, using both PTH and PTHrP.

We first compared the ability of PTH1R to activate different G proteins, using BRET-based biosensors^38^, which respond to PTH1R activation with a decrease in BRET between their Gγ subunit labelled with the bioluminescent donor NanoLuc and their Gα subunits tagged with the acceptor cpVenus (**Fig. 5**). HEK293 cells were transiently transfected with the specific BRET biosensor along with PTH1R_wt_ with or without RAMP2_wt_ as described above. Experiments were conducted in microtiter plates and BRET signals were recorded over time until they reached their maximal response.

Significant PTH-induced changes in BRET were observed with all four G protein biosensors. They were all concentration-dependent with EC_50_-values in a range reflecting G protein preferences of this receptor in order G_s_ > G_q_ > G_13_ > G_i3_ (*SI Appendix*, Fig. S6 upper panels; *SI Appendix*, Table S1). Two changes in the G protein activation patterns were notable. First, the presence of RAMP2_wt_ caused a more rapid activation of Gs by PTH with a brief initial overshoot peaking at 2 min after receptor activation (**Fig. 4*A***); when measured at this time point, the presence of RAMP2_wt_ increased the PTH-induced BRET change (**Fig. 4*A***). Second, RAMP2_wt_ caused a specific increase in potency for PTH-triggered G_i3_ activation, resulting in a major difference of Gi3 activation by low concentrations of PTH (**Fig. 4*B***, SI Appendix, Fig. S6 G; *SI Appendix*, Table S1); at 10 nM PTH, the presence of RAMP2_wt_ markedly accelerated Gi3 activation (**Fig. 4*B***). In contrast, the potencies, and efficacies of PTH-stimulated activation for G_q_ and G_13_ were not affected by RAMP2 (*SI Appendix*, Fig. S6) (upper panels; *SI Appendix*, Table S1).

**Fig. 4.**
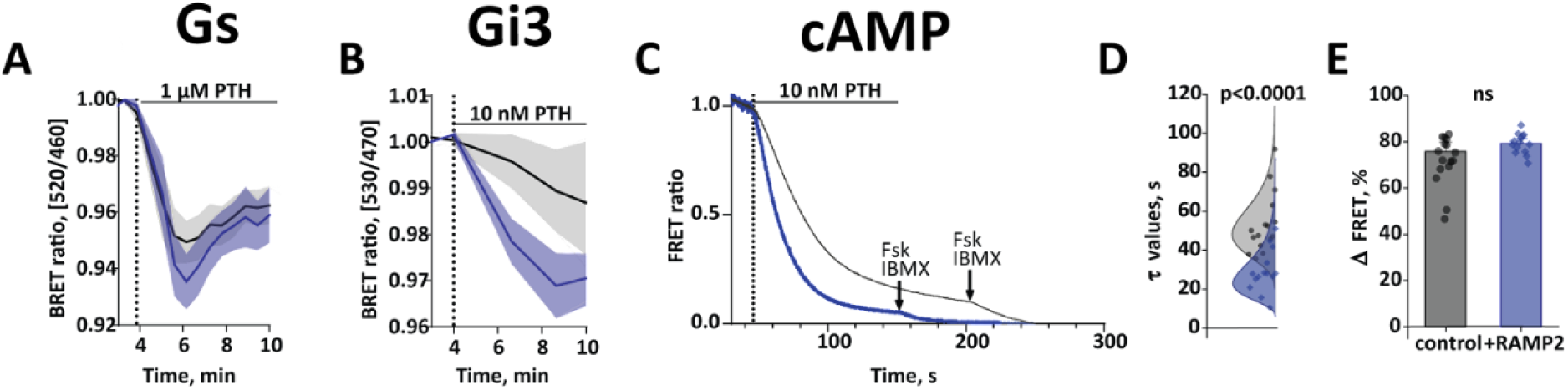
RAMP2 effects on PTH-stimulated G protein activation and cAMP accumulation. *(A, B)* HEK293 cells transiently transfected with cDNA encoding for BRET biosensors of G proteins: G_s_ (*A*) and G_i3_ (B) along with PTH1R_wt_, with or without RAMP2_wt._ BRET signals were recorded in a plate reader from cells stimulated with indicated concentrations of PTH. Shown are time courses of agonist stimulation. Data are means ± SEM of n = 3 independent experiments performed in duplicates or more. For further statistics and concentration response curves see *SI Appendix*, Fig. S6, Table S1 and S2. (*C*) HEK293 cells transiently transfected with cDNA encoding for the cAMP-based FRET biosensor (Epac-S^H187^), along with PTH1R_wt,_ with or without RAMP2_wt_. Shown are representative ratio traces of PTH-mediated FRET changes in single HEK293 cells, recorded in a microscopic FRET setup. Horizontal lines indicate application of 10 nM PTH with a rapid superfusion system. The arrow indicates addition of 10 µM Forskolin and 100 µM IBMX after signal saturation. Traces were normalized to the baseline (set to 1) and plateau after stimulation with Forskolin and IBMX (set to 0). Traces are representative of n = 16 cells (control) and n = 14 cells (+RAMP2_wt_), acquired in two independent experiments. (*D*) Time constants *τ* of PTH-induced FRET changes derived from traces as in panel *C*, calculated from mono-exponential curve fitting. The data were fitted with a lognormal distribution. Median value and 95% confidence interval (CI) were: PTH1R_FRET_ = 49 ms [45, 62 CI], n = 14 cells; RAMP2_SNAP_ = 28 ms [23, 35 CI], n = 14 cells. A Mann-Whitney test was used to assess a significant difference between the groups (p < 0.001). (*E*) Effects of RAMP2 on the amplitude of the FRET signals induced by PTH. Bars represent means ± SEM in % of the ΔFRET amplitudes from all cells examined: Epac-S^H187^+PTH1wt = 75.8 ± 4.1 % (n = 16 cells); Epac-S^H187^+PTH1wt +RAMP2_wt_ = 79.3 ± 2.1 %, (n = 14 cells). A t-test was used to assess a significant difference between the groups (ns, p > 0.05).

PTHrP elicited similar decreases in BRET for all G protein biosensors (*SI Appendix*, Fig. S6, lower panels), which occurred with similar time courses as for PTH. However, in contrast to PTH, the co-expression of RAMP2_wt_ did not significantly change the potency or efficacy of PTHrP-stimulation for any of the G proteins analysed (*SI Appendix*, Fig. S6 lower panels and Table S2).

To assess whether downstream effects corresponded to those seen at the G protein level, we also measured cAMP accumulation, using Epac-S^H187^ cAMP biosensor^39^ (**Fig. 3**). The amplitude of cAMP accumulation recorded in microtiter plates was similar in RAMP2_wt_ expressing cells and in control cells at all concentrations of PTH (**Fig. 4*E***, SI Appendix, Fig. S6I). Strikingly, in line with the accelerated G_s_ activation, the speed of cAMP accumulation measured at a single cell level with a rapid superfusion system^11^, in agreement with results obtained with the PTH1R_FRET_ biosensor, were affected by RAMP2: in particular, RAMP2 accelerated the PTH-induced cAMP accumulation (**Fig. 4*C-E***). The time constant (***τ***) of activation was decreased from median time constant 45 s to 25 s by the presence of RAMP2 (**Fig. 4*D***).

### RAMP2 effects on non-G protein signalling

In addition to activation of G proteins, agonist-activated PTH1R is phosphorylated by G protein-coupled receptor kinases (GRKs) and then binds β-arrestins, thereby triggering receptor internalization and signalling by extracellular signal-regulated kinases (ERKs). The latter process appears to have different conformational requirements compared to G protein activation^21-23,40^. We therefore set out to also assess the effects of RAMP2 on these signalling mechanisms, employing various BRET and FRET biosensors to quantify GRK2 and β-arrestin2 recruitment to the PTH1R along with ERK activation. To monitor recruitment of GRK2 and or β-arrestin2, we used BRET assays, in which PTH1R was tagged with the donor NanoLuc (PTH1RNanoLuc), while GRK2 and β-arrestin2 were tagged with YFP and mVenus respectively^4^. We measured BRET signals at the time of the maximal response after full agonist occupancy (**Fig. 5*A,C,E***).

**Fig. 5.**
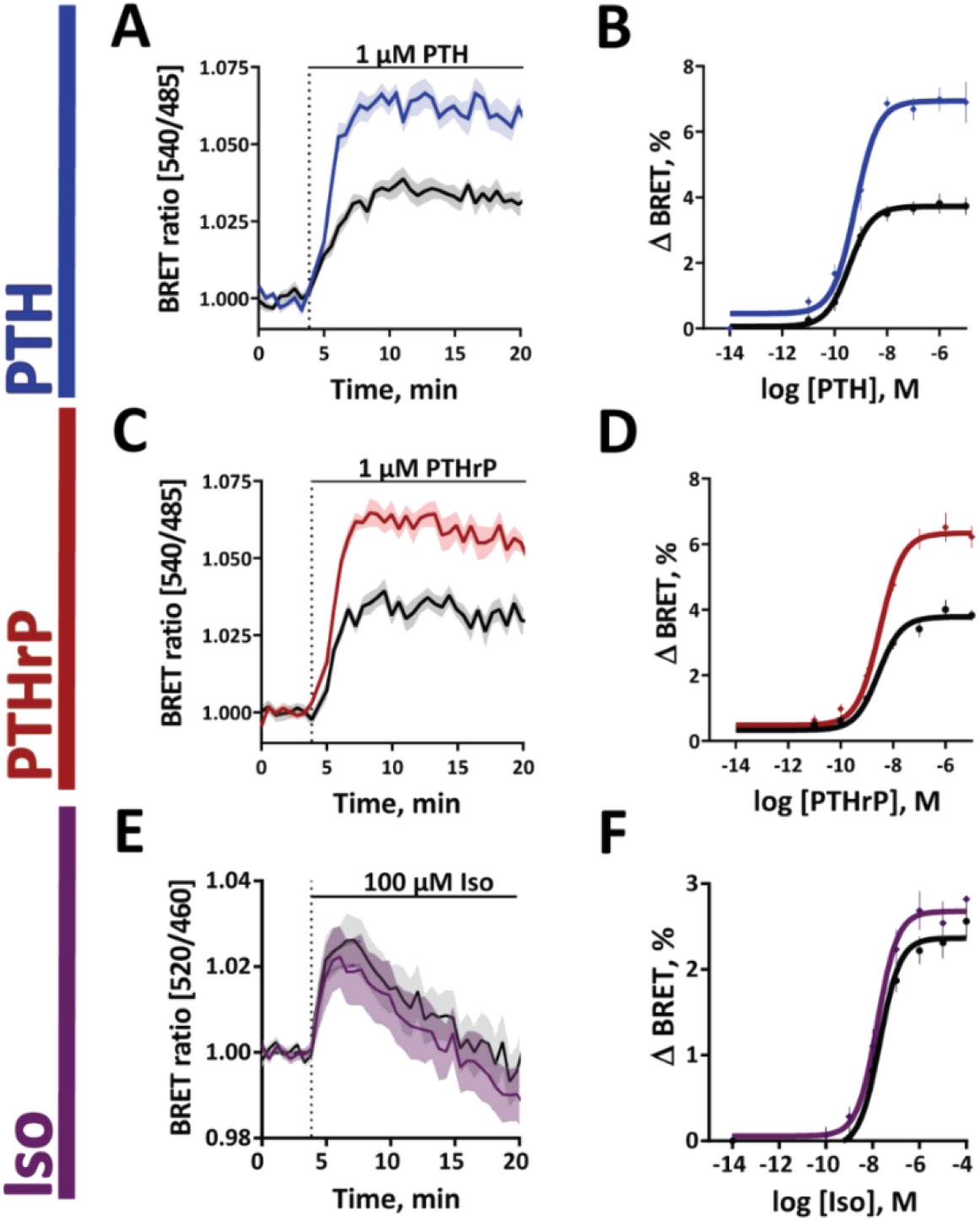
RAMP2 effects on β-arrestin recruitment. HEK293 cells were transiently transfected with cDNA encoding for: β-arrestin2_mVenus_ along with PTH1R_NanoLuc_ (*A - D*) or β2AR_NanoLuc_ (*E, F*), with or without RAMP2_wt_. BRET signals were recorded in a plate reader from cells stimulated with PTH (*A, B*), PTHrP (*C, D*) or isoprenaline (*E, F*). Shown are averaged time courses of agonist stimulation (*A, C, E*) and corresponding concentration-response curves (*B, D, F*), fitted with a three-parameter concentration-response curve fit. Data are means ± SEM of at least n = 3 independent experiments performed in quadruplicates or more. For further statistics and results see *SI Appendix*, Table S3 and S4.

The main change induced by RAMP2 in this series of experiments was a marked increase in β-arrestin2 recruitment; visualized as a significant increase in amplitude of BRET ratio for both PTH and PTHrP (**Fig. 5*A,C***, Table S3, S4). This increase was visible at all concentrations of PTH and PTHrP (**Fig. 5*B,D***). Control experiments showed that RAMP2 did not affect β-arrestin2 recruitment in the absence of receptor stimulation (*SI Appendix*, Fig.S7L). Further control experiments indicated the specificity of the effects of RAMP2, because it did not alter β-arrestin2 recruitment to the β2-adrenergic receptor, (**Fig. 5*E,F***, Table S5), which does not interact with RAMP2^29,33^ (**Fig. 1*C***).

In contrast to these major and very robust effects on β-arrestin2 recruitment, there were only minor or no effects on GRK2 recruitment and on ERK activation (*SI Appendix*, Fig. S8, Table S3, S4).

Taking all data on PTH1R signalling together, we demonstrate two major effects of RAMP2: 1) a PTH-selective increase in the speed of stimulating Gs and potency of G_i3_ activation (**Fig. 4**; *SI Appendix*, Fig. S6), 2) an increase in β-arrestin2 recruitment which is seen for both agonists (**Fig. 5)**. Interestingly, the latter effects are not translated into increased ERK signalling by PTH1R. However, the speed of the increased recruitment of β-arrestin2 corresponds to the kinetics of the overshoot in Gs activation, in line with the role of β-arrestins to limit G protein activation.

### Structural models of putative PTH1R-ligand-RAMP2-Gs complexes

The modelling approach carried out here resulted in two different proposals for the complex formation of RAMP2 with PTH1R and PTH ligand variants (*SI Appendix*, Fig.S12A). In model version I, the RAMP2-ECD is bound to the PTH1R-ECD, as suggested by the known CLR-CGRP-RAMP1 complexes^51^, but the overall receptor-ECD orientation to the TM region is maintained according to the known PTH1R-LA-PTH-Gs complex^16^. In this scenario - with unchanged the complex arrangement and ligand conformation, but additionally bound RAMP2 - no significant changes in ligand binding or inter- or intramolecular interactions are apparent despite few receptor-RAMP2 interactions mainly in the TM region.

In contrast, taking the previously solved CLR-CGRP-RAMP1-Gs complex^51^ as a structural template for modelling a putative complex between PTH1R-PTH/PTHrP-Gs with RAMP2 (*SI Appendix*, Fig. S12B), several highly relevant structural parts are potentially altered with modified interaction patterns both intra- and intermolecular compared to the known PTH1R-LA-PTH complex^16^. The linker region of RAMP2 (e.g. F138, D140) would interact with the C-terminus of PTH1R EL2, in addition to specific RAMP2-ECD to PTH1R-ECD contacts (e.g. receptor-RAMP2: Q45-R97). The PTH1R ECD can contact EL3 (e.g. E431), which is expected to affect directly neighboring helices TM6 and TM7, which are known to be important for signal transduction and regulation. Furthermore, the ligand-receptor interaction pattern is modified in this model (**Fig. 6**), assuming additional contacts (e.g. receptor-PTH: D133-K13).

**Fig. 6.**
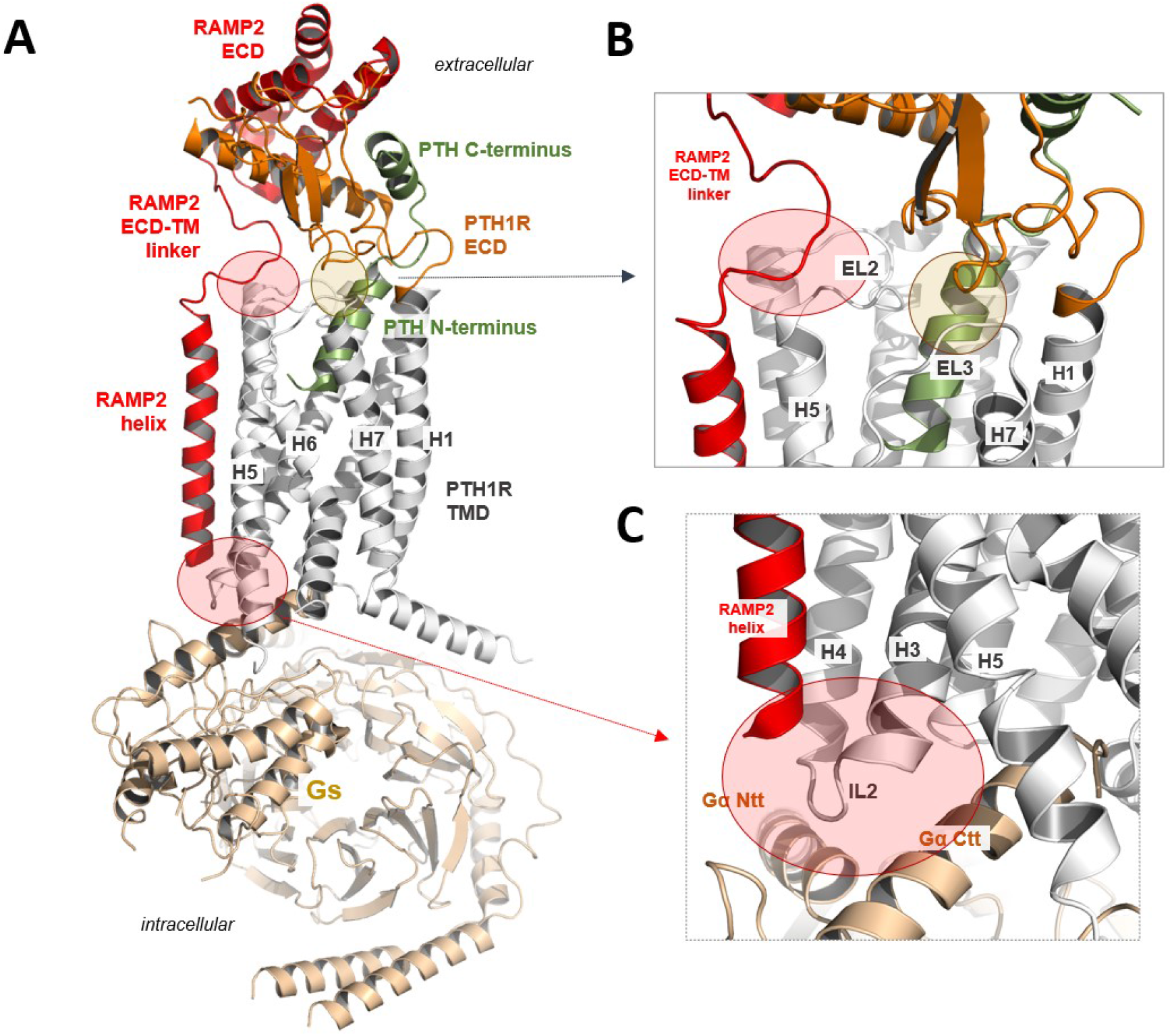
Putative RAMP2 binding mode in a PTH1R-PTH-Gs complex model. (*A*) A homology model between PTH1R-PTH-Gs and RAMP2 suggests several specific contacts between the components of this complex. RAMP2 should interact at the extracellular region with the EL2 of the receptor (*B*), but also at the intracellular site with the IL2 and adjacent transitions to receptor helices 3 and 4 (*C*) (red translucent circles), a region that is associated with G-protein binding (Ntt, N-terminal tail; Ctt, C-terminal tail). There are also new intramolecular contacts from the PTH1R ECD to the TMD (transparent green circle), which are not observable in the recently determined PTH1R complex structure^16^ (PDB ID: 6nbh). In this model the receptor ECD would interact directly with the EL3, but potentially also with EL1. In addition, several new receptor ECD-ligand contacts are feasible, such as K13 (PTH) and D133 of the receptor.

## Discussion

RAMPs have co-evolved and are co-expressed with several GPCRs^28,33,44,45^. More than 40 partner proteins have been described to interact with them^29,46-48^. The effects of such interactions have been studied in most detail for their prototypical interaction partners – the class B GPCRs calcitonin and calcitonin-like receptors (CTR, CTRL). RAMP’s binding to these receptors has been described to facilitate transport of the receptors to the cell surface, to change ligand specificity, and also to alter their downstream signalling cascades^49^. However, little is known about how RAMPs might affect the activation process of a GPCR itself.

Here, we investigated possible modulatory effects using the class B PTH1R as a model system. Among the three RAMPs, this receptor showed a clear preference for RAMP2. Co-expression of RAMP2 with PTH1R increased the activation speed of the receptor several-fold, and it reduced the amplitude of the activation signal by ≈2-fold. Both effects were seen with a FRET-sensor as well as a new cpGFP-based biosensor with similar effect sizes. In line with the smaller signal amplitude, basal FRET of the FRET biosensor was decreased by RAMP2, i.e. altered in a way similar to the effects of (partial) agonists. This might suggest that RAMP2 induced a partially pre-activated state, from which agonist-induced activation may proceed with much greater speed. Based on the various fluorescence and FRET readouts, this pre-activated state appears to be distinct both from the inactive and from the fully active state. Given that RAMP2 changed the basal and stimulated conformations and did not have an impact on the basal FRET ratio of G protein biosensors or other downstream effectors, we conclude that RAMP2 appears to induce unique receptor conformations.

Interestingly, these effects were ligand-specific: they were prominent for PTH, but essentially absent for PTHrP. This was true for both, the increase in activation speed and the decrease in amplitude of the activation signals. It suggests, first, that the two endogenous ligands can be regulated in a differential manner and, second, that RAMP2 can exert very specific and subtle effects on PTH1R. Such specific conformational changes are somewhat reminiscent of analogous kinetic effects that have been observed in homo- and hetero-dimeric GPCRs^13,15^. Ligand-dependent effects were previously described for some with RAMPs interacting GPCRs^50,51^.

RAMP1-CTRL hetero-oligomers have been shown to RAMP-specifically propagate extracellular dynamics to the cell interior and through that to control receptor phenotypes^52^. Structurally, all RAMPs are tightly packed with their interacting GPCR partner, being placed between transmembrane domains 3, 4, and 5 and making contacts with the second extracellular loop (EL2)^51,52^. Thus, RAMPs are placed near structural motifs that are governing GPCR activation.

In an attempt to interpret our data in a structural manner, we performed structural homology modelling of PTH1R-ligand-RAMP2-Gs complexes to suppose potential intermolecular interactions, pre-activation and possible impact on the intracellular signalling of PTH1R-RAMP2 oligomer (**Fig. 6*A***). Starting from available structural information^16,51,52^ *(SI Appendix*, Figures S10-S12), our model provides indications that RAMP2 mediated pre-activation might originate from the interaction of <u>the RAMP2-linker</u> (**Fig. 6*B***) with the receptor EL2 (upper part of TM5) and ECD (red circle), which is in contact with the EL3 (green circle) that connects TM6 and TM7. The extracellular contributions of new intra- and intermolecular contacts might cause pre-activation or stabilize a pre-activated conformation of the receptor, comparable to scenarios known from several GPCRs with activated “intramolecular agonists”^53^. In this case, the binding of RAMP2 would act as an initial (partial) activation trigger. Moreover, the EL2 of PTH1R has been recognized as an allosteric hotspot, where selective modulation or mutational alterations can affect the balance between G protein coupling and β-arrestin-driven signalling^54,55^. In addition, the <u>RAMP2-helix</u> contacts the receptor-IL2 (**Fig. 6*C***, red circle) and the adjacent connections to TM3 and TM4 intracellularly. This receptor part is highly interrelated with G protein binding and thus any modification by RAMP2 binding should result in altered functional receptor properties even in the basal state. IL2 has been shown to predispose to constitutive receptor activation in several class A GPCRs, as demonstrated by mutagenesis studies^56,57^ and may also be highly relevant in the class B.

In line with the predicted altered G protein function we observed a number of remarkably specific changes on PTH1R downstream signalling by RAMP2. First, we found that RAMP2 caused a specific and selective increase in G_s_ and G_i3_ activation kinetics by PTH, suggesting selectivity in the modulation of G protein coupling. Again, this effect was ligand specific, being much more pronounced for PTH than for PTHrP. Among the non-G protein interactions of the PTH1R, we observed a substantial and specific increase on β- arrestin2 recruitment by RAMP2; this recruitment paralleled the overshoot in Gs activation, suggesting that it might limit Gs activation and signalling. In contrast, increased β-arrestin2 recruitment was not translated to other β-arrestin-dependent cascades such as ERK activation. It remains to be seen whether the increased β-arrestin2 recruitment BRET signal is due to an increased amount of β-arrestin2 recruited, or to a different state of β-arrestin2 induced by the PTH1R/RAMP complex compared to PTH1R alone^58,59^, and whether β-arrestins are somehow shielded from their downstream interaction partners by RAMP2. Since specific β-arrestin-dependent transcription programs have been identified for the PTH1R, which might provide a therapeutically interesting pathway to increase bone mass^22,60^, RAMP2-dependent modulation of these pathways may provide a new type of pharmacological target. This mechanism might be pharmacologically attractive since PTH1R/RAMP2 is not an obligate but a tissue-dependent hetero-dimer, which might represent a source of unique, tissue-dependent biased signalling patterns in different tissues and, thus, a unique pharmacological targeting approach^49,60^. The development of erenumab, an antibody that specifically targets the CTRL/RAMP interface indicates that such a GPCR/RAMP interface can be exploited as a pharmacological target^61^.

In summary, our data, together with recent structural insights, highlight a new unique conformation of the PTH1R when interacting with its regulator RAMP2. This presumably pre-activated state promotes faster and ligand-specific activation of PTH1R and controls its signalling specificity. These data illustrate a critical role of a RAMP in GPCR activation and signalling.

## Materials and Methods

### Chemicals

The peptide ligands parathyroid hormone PTH (1-34) (human, H-4835-GMP, #4033364) and parathyroid hormone-related peptide (PTHrP) (1-34) (human, mouse, rat; #4017147) were from Bachem. SNAP-Cell 647-SiR (#S9102S) was from New England Biosciences. Anti HA-tag antibody (ab9110) was from Abcam and anti-rabbit IgG, HRP-linked antibody (#7074P2) was from Cell Signalling. 3,3′,5,5′-Tetramethylbenzidine (TMB, #T8665) was from Sigma-Aldrich. NanoBRET™ Nano-Glo® Substrate - furimazine (#N1663) and HaloTag® NanoBRET™ 618 Ligand (#G9801) were from Promega. Dimethylsulfoxide (DMSO, #A994.2) for cell culture was from Carl Roth GmbH & Co. KG. Bovine Serum Albumin (BSA, #SAFSA7030) was from VWR International.

### Molecular cloning

All PTH1R-based constructs were cloned from human full-length PTH1R. Plasmids were either created by molecular restriction cloning or by the Gibson Assembly technique (New England Biolabs, Inc.). HA-PTH1R_mTurquoise/mCitrine_ (PTH1R_FRET_) and HA-PTH1R_mCitrine_ were modified from previously described biosensors^11,34^. For HA-PTH1R_NanoLuc_, NanoLuc was fused to the C-terminal of HA-PTH1R_wt_^60^.

PTH1R_cpGFP_ biosensor was cloned into pCMV Twist vector and designed according to the previously described dLight1 cpGFP biosensor^36^ and synthesized by Twist Bioscience, California, USA. Influenza A signalling peptide (MKTIIALSYIFCLVFADYKDDDDA) was fused to the N-terminus of PTH1R and LSSLI-cpGFP-NHDQL was inserted between Lys388 and Arg400 in the third intracellular loop.

Wildtype RAMP constructs were a gift from Annette Beck-Sickinger (University of Leipzig, Leipzig, Germany). RAMP_mCitrine_ and RAMP_SNAP_ were generated by fusing mCitrine or SNAP-tag to the C-termini of RAMPs. SNAP-tag sequence was amplified from a SNAP-GABA_B1_ receptor template, kindly provided by Jean-Philippe Pin (Institut de Génomique Fonctionnelle, Montpellier, France). The C-terminus of the CAAX sequence was tagged with mCitrine or mTurquoise2. Nuclear EKAR (Cerulean-Venus) was a gift from Karel Svoboda (Janelia Research Campus, Virginia, USA; Addgene plasmid #18682), biosensor Epac-S^H187^ was a gift from Kees Jalink (The Netherlands Cancer Institute, Amsterdam, Netherlands), GRK2_YFP_^63^ was described previously. For β_2_AR_mCitrine_ and β_2_AR_NanoLuc_, mCitrine and NanoLuc were fused to the C-terminus of β_2_AR, respectively. β-arrestin2_mVenus_ was modified from previously described β-arrestin2_EYFP_^65^ by exchanging EYFP for mVenus.

The expression vector in all plasmids was pcDNA3(+) unless otherwise noted.

All constructs were verified by sequencing by Eurofins or LGC genomics. All plasmids and sequences are available from the authors upon request.

### Cell culture

Different clones of human embryonic kidney cells (HEK293) were employed. HEK293 (ECACC #85120602, CRL-1573, ATCC) was used for the generation of a stable cell line, HEK293T for most experiments (ECACC #96121229, Sigma-Aldrich) and HEK293A (R70507, Thermo Fisher) for plate reader experiments of G protein activation and internalization. To all three clones of HEK293 cells we refer as HEK293 in the main section.

Cells were grown in Dulbecco’s Modified Eagle’s Medium (DMEM, Pan Biotech) supplemented with 2 mM L-glutamine (Pan Biotech), 10 % fetal calf serum (Biochrome), 100 μg/mL streptomycin and 100 U/mL penicillin (Gibco), at 37°C with 5 % CO_2_. Cells were washed with phosphate-buffered saline (PBS, Sigma-Aldrich) and passaged with 0.05 %/0.02 % Trypsin/EDTA (Pan Biotech) every 2-3 days when reaching 80 % confluency. Cells were routinely tested for mycoplasma infection using MycoAlert^™^ Mycoplasma Detection Kit (Lonza). Cells were not contaminated with mycoplasma.

### Creation of stable cell line

HEK293 cells were used to develop a stable cell line of PTH1R_FRET_ biosensor. Cells seeded into 100 mm dishes were transfected at a confluence of 60 % with 2 µg of cDNA encoding PTH1R_FRET_ with Lipofectamine 3000 Transfection Reagent Kit (Qiagen), according to the manufacturer’s protocol. Transfected clones were selected with 600 μg/mL G-418 (VWR International) and sorted with a flow cytometer. Monoclonal single clones were grown in DMEM supplemented with 200 μg/mL G-418. For further experiments, the best clone was selected based on the brightness and amplitude of the saturating PTH stimulation of PTH1R_FRET_ biosensor in plate reader experiments.

### Seeding and transfection

Coverslips or microtiter plates were covered with poly-D-lysine (PDL) for 30 min, washed 2x with PBS and left to dry before seeding. For microscopy experiments, 2×10^5^ cells were seeded onto 25 mm coverslips (Sigma-Aldrich) into a 6-well plate. After 24 h cells were transfected with Lipofectamine 3000 (Qiagen), according to the manufacturer’s protocol. For all transfections, the ratio of PTH1R:pcDNA3/RAMP was 1:1, unless otherwise noted. The empty backbone of pcDNA3 was used throughout to maintain a consistent level of total cDNA. For plate reader experiments, each methods section contains a detailed description of seeding and transfection protocol.

### FRET acceptor photobleaching (FRET-AB) in confocal microscopy

Cells were imaged 36 h after transfection. Coverslips were mounted onto Attofluor™ chamber (Fisher Scientific) and washed once with FRET buffer (137 mM NaCl, 5 mM KCl, 1 mM CaCl2, 1 mM MgCl2, 20 mM HEPES, pH 7.4) containing 0.1 % (wt/vol) BSA (AppliChem). Cells were kept in FRET buffer at room temperature throughout the experiment.

The chamber was mounted onto a Leica SP8 confocal laser-scanning microscope, equipped with an oil immersion objective (HC PL APO CS2 40x/1.3 NA). LAS X microscope control software and the Leica FRET-AB wizard tool were used to perform experiments. A 1.5 mW white light laser was set to 1 % and a 431 nm laser line was used at 1 % power for donor imaging. For acceptor imaging, a 512 nm laser line at 1 % power was used and for the bleaching step increased to 100 % for 10 frames. 512 × 512-pixel images were acquired with a hybrid detector in standard mode. Emission of donor channel was recorded within 440 – 512 nm and emission of acceptor channel was recorded within 517 – 620 nm. The zoom factor was set to 5.5 x, resulting in a pixel size of 103 nm, and the laser scanning speed was set to 400 Hz. Fixed-size regions of interest (ROI) were selected on the membrane of the cell. For intramolecular FRET-AB experiments, ROIs expressing both PTH1R_FRET_ and RAMP2_SNAP_ were selected.

FRET efficiencies were calculated with the manufacturer’s Wizard tool, based on the provided Equation 1, where I denotes the fluorescence emission intensity:

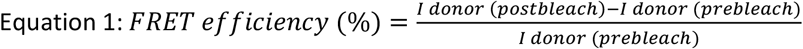

A maximum of 4 cells was taken for analysis per image. To ensure co-expression integrity and enough bleaching of the acceptor only cells with initial emission ratios (mCitrine/mTurquoise2) within 0.25 and 4, and bleaching > 20 %, were considered for statistical analysis.

### SNAP-tag labelling

Before FRET experiments, coverslips, expressing a combination of PTH1R_FRET_ and RAMP2_SNAP_ were labeled with 1 µM SNAP-Cell 647-SiR in serum-free Fluorobrite^™^ DMEM (Gibco) for 30 min and kept in the incubator at 37°C and 5 % CO_2_. Excessive dye was washed by exchanging medium 3 times every 10 min.

### Single-cell PTH1R biosensor experiments in intact cells

Cells were imaged 36 h after the transfection. Coverslips were mounted onto an Attofluor™ chamber and washed once with FRET buffer. Cells were kept in FRET buffer at room temperature throughout the experiment. The chamber was mounted onto an inverted microscope (DMi8, Leica Microsystems, Germany), equipped with an oil-immersion objective (HC PL APO 63×/1.40-0.60 oil, Leica Microsystems), dichroic beamsplitter T505lpxr (Visitron Systems, Puchheim, Germany), and Xenon lamp coupled with a continuously tunable Visichrome high-speed polychromator (Visitron Systems). Images were acquired with an sCMOS camera (Prime 95B, Teledyne Photometrics, USA) using a dual image splitter (OptoSplit II, Cairn Research, UK). Image sequences had 40 ms (PTH1R_FRET_) or 100 ms (Epac-S^H187^) acquisition intervals and were recorded with the VisiView 4.0 software (Visitron Systems). Ligand application was performed using a solenoid valves perfusion system with a 200 µm inner diameter manifold-tip (Octaflow II, ALA Scientific Instruments, USA).

To check for RAMP2_SNAP_-expressing cells, cells were excited at 640 nm for 100 ms and fluorescence emission was recorded at 690/50 nm.

For FRET experiments, cells expressing PTH1R_FRET_ or Epac-S^H187^ were excited with 445 nm, and fluorescence emission was simultaneously recorded at 470/24 nm and 535/30 nm. Cells expressing PTH1R_cpGFP_ were excited at 483 nm and fluorescence emission was recorded at 506 nm.

### Fluorescence spectrum and fluorescence experiments in the plate reader

3×10^6^ HEK293T cells were seeded into 100 mm dishes and transfected after 24 h with the combination of PTH1R_FRET_/PTH1R_cpGFP_ and pcDNA3/RAMP2_wt_/RAMP2_SNAP_ or PTH1R_wt_, Epac-S^H187^/EKAR biosensor, and pcDNA3/RAMP2_wt_/RAMP2_SNAP_. Combinations were transfected at a ratio of 1:1 or 1:1:1, respectively. 24 h after the transfection, cells were transferred to PDL-precoated black-wall, black-bottomed 96-well plates (Brand) at a density of 70,000 cells/well. 36 h after the transfection, cells expressing biosensors were washed and medium was substituted with FRET buffer. Plate reader experiments were conducted at 37 °C using a Synergy Neo2 plate reader (BioTek, Vermont, USA) equipped with a monochromator and filter optics. 10 excitation flashes were applied per data point.

For the fluorescence emission spectrum of PTH1R_cpGFP_, cells were excited at 455/10 nm and fluorescence emission was recorded with 1 nm resolution within 500 -660 nm.

For amplitude experiments and generation of concentration-response curves, basal reads for 5 min were recorded in 90 µL FRET buffer. Subsequently, 10 µL of 10-fold ligand solution or FRET buffer was applied to each well and the stimulated reads were recorded for a further 10 min. For FRET experiments, 420/50 nm excitation filter and 485/20 nm and 540/25 nm dual emission filter were used. For PTH1R_cpGFP_-expressing cells, 485/20 nm excitation and 516/20 nm emission filters were used.

For PTH1R_FRET_, EKAR or Epac-S^H187^ biosensor-expressing cells, expression levels were measured with monochromator optics. Cells were excited at 510/20 nm and fluorescence emission was recorded at 560/20 nm.

### Live-cell ELISA

3×10^6^ HEK293T cells were seeded into 100 mm dishes and transfected 24 h later with a combination of PTH1R_FRET_ and pcDNA3/RAMP2_wt_/RAMP2_SNAP_ or PTH1R_cpGFP_ and pcDNA3 (no HA-tag control) at a ratio of 1:1. The medium was exchanged after 12 h and 24 h after the transfection and the cells were transferred to PDL-precoated transparent 96-well plates (Brand) at a density of 70,000 cells/well. 48 h later, cells were washed 2x with 0.5 % BSA/PBS. Subsequently, cells were incubated for 1 h at 4 °C with rabbit anti-HA tag antibody (1:1000) in 1 % BSA/PBS. Following incubation, cells were washed 4x with 0.5 % BSA/PBS and incubated with goat anti-rabbit IgG, HRP-linked antibody (1:4000) in 1 % BSA/PBS for 1 h at 4 °C. Finally, cells were washed 3x with 0.5 % BSA/PBS, and 50 µl of the peroxidase substrate TMB was added. Following a 30 min incubation and development of a blue product, absorbance was recorded at 665 nm using a Neo2 plate reader.

### BRET-based G protein activation assay

HEK293A cells were transfected with PTH1R_wt_, G protein BRET biosensor, and pcDNA3/RAMP2_wt_ at a ratio of 1:1:1. Constructs were transfected in suspension with Lipofectamine 2000 (2 µl transfection reagent/1 µg total cDNA) according to the manufacturer’s protocol and seeded into a PDL-precoated, white-wall, white-bottomed 96-well microtiter plate (30,000 cells/well). 48 h after the transfection, cells were washed with Hanks Balanced Salt Solution (HBSS) and incubated with 90 µL of a 1:1000 (vol:vol) stock solution of furimazine in HBSS. 5 min later, 3 consecutive reads were recorded as basal reads. Subsequently, 10 µL of a 10-fold ligand solution or HBSS was applied to each well and the stimulated reads were recorded.

All experiments were conducted using a CLARIOstar plate reader (BMG Labtech, Ortenberg, Germany) recording NanoLuc and cpVenus emission with 450/80 nm (gain 3600) and 530/30 nm (gain 4000) monochromator settings, respectively, and an integration time of 0.3 s.

### BRET-based GRK2 recruitment, β-arrestin2 recruitment, and Gs protein activation assay

HEK293T cells were transfected with GRK2_EYFP_, PTH1R_NanoLuc_ and pcDNA3/RAMP2_wt_; β-arrestin2_mVenus_, PTH1R_NanoLuc_ and pcDNA3/RAMP2_wt_ or Gs protein BRET biosensor, PTH1R_wt_ and pcDNA3/RAMP2_wt_ at a of ratio 1:1:1. Combinations were transfected with Lipofectamine 3000 according to the manufacturer’s protocol with a total of 6 µg of cDNA. After 12 h medium was exchanged and after 24 h cells were transferred into a white-wall, white-bottomed 96-well microtiter plate, at a density of 60,000 cells/well. 24 h after the reseeding, the medium was removed, and cells were washed once with FRET buffer and incubated with 90 μL of a 1:1000 (vol:vol) stock solution of furimazine in FRET buffer. 5 min later, basal reads were recorded for 4 min and subsequently, 10 µL of 10-fold ligand solution or FRET buffer was applied to each well and the stimulated reads were further recorded.

Measurements were performed at 37 °C using a Synergy Neo2 Plate Reader with the NanoBRET filter set, integration time per data point was set to 0.3 s and gain to 100/120 (GRK2 recruitment) or 90/110 (β-arrestin2 recruitment, Gs activation).

GRK2_YFP_- and βarr2_mVenus_-expressing cells were excited at 510/20 nm and fluorescence emission was recorded at 560/20 nm for quantification of expression level.

### Data analysis and statistics

For microscopic FRET experiments, fluorescence emission time courses of both FRET donor and acceptor were routinely corrected for background and spectral bleedthrough, and the FRET ratio was calculated as described earlier^9,64^. For calculating τ, agonist-independent changes in FRET due to photobleaching were subtracted. The decrease in FRET ratio was fitted to the one-phase decay equation: r(t) = A × (1 − e^−t/τ^) (*Equation 2*), where τ is the time constant (s) and A is the amplitude. X_0_ was constrained to the time when the decay began. Δ FRET values were calculated as normalized differences between basal and stimulated FRET ratios.

For plate reader experiments, the data were analyzed in Microsoft Excel and if needed, wells out of the fluorescence range of plate readers were excluded as an outlier. For FRET and BRET experiments, raw RET ratios were defined as acceptor emission/donor emission. RET ratios before ligand/buffer addition were averaged and defined as RET_basal_. To quantify ligand-induced RET changes, ΔRET was calculated for each well and time point as percent over basal ([(RET_stim_− RET_basal_)/RET_basal_] × 100) (*Equation 3*). Subsequently, the average ΔRET of buffer-treated control wells was subtracted. To reduce the fluctuation of the BRET ratio, three consecutive BRET ratios were averaged before and after ligand addition^66^. Concentration-response curve experiments were fitted using a three or four-parameter logistic curve fit as stated in corresponding figure legends.

Statistical differences were evaluated using a one-way ANOVA test followed by Tukey multiple comparisons, Brown-Forsythe ANOVA, followed by Dunnett T3’s multiple comparisons test, Student’s t-test, Mann Whitney test, or extra-sum-of squares F-test. Each figure legend contains a description of statistical treatment. Differences were considered significant for values of p < 0.05. The data were analyzed and visualized using Microsoft Excel 2016 (Microsoft, Washington, USA), GraphPad Prism software 8.1.2 (GraphPad Software, California, USA), and OriginPro 2018 software (OriginLab, Massachusetts, USA).

## Acknowledgments

We thank all members of the Lohse lab, especially Ali Isbilir and Jan Möller, for valuable discussions on the manuscript. G.K. and P.S. were supported by the DFG - German Research Foundation through CRC 1423, project number 421152132, subprojects A01/A05/Z03; through CRC 1365, project number 394046635, subproject A03; through Germany’s Excellence Strategies − EXC2008/1 (UniSysCat) – 390540038, and two European Union’s Horizon 2020 MSCA Program under grant agreement 956314 [ALLODD]. This work was supported by the German Research Foundation (SFB688-B08, 427840891) and Elitenetzwerk Bayern, Receptor Dynamics program and Federal Ministry of Research (BMBF; 03V0830) to Martin. J. Lohse. and University of Nottingham Anne McLaren Research Fellowship to Isabella Maiellaro.

