## Supplementary Information for "Functional modulation of PTH1R activation and signalling by RAMP2"

### Supplementary methods

**Structural homology modelling of PTH1R-ligand-RAMP2-Gs complexes**

To obtain three-dimensional homology models of putative arrangements between RAMP2 and PTH1R bound with the peptidic ligands PTHrP or PTH, several available structural templates were used[16,51].

It should be noted that the native PTH ligand in complex with the PTH1R receptor and both together with a cofactor protein RAMP have not yet been structurally determined. The following aspects should be mentioned when building a complex RAMP2-PTH1R-PTH structural model.

First, the PTH1R structure determined so far in complex with long-acting PTH and Gs-protein (LA-PTH, Fig.S10 A-B1) provides a suitable receptor starting template for the design of complexes with both PTH and PTHrP as well as RAMP.

Second, an appropriate structural template for detailed RAMP binding to PTH1R is the CLR-CGRP complex, which has also already been solved (Fig. S10 B2). This complex shows an overall and detailed binding mode of RAMP1 and provides information about the orientation of the membrane-spanning helix (RAMP-helix) and the extracellular domain (RAMP-ECD) of the RAMP1 protein fold at the receptor-ligand complex. A RAMP2-adrenomedullin-receptor structure (AM1R) is also available[52] (PDB ID: 6uun), but most of the side chains in the extracellular parts are not visible in the structure, making it less suitable for homology modeling purposes.

Third, it is further important to note that the spatial orientation of the receptor extracellular domain (receptor-ECD) in complex with the ligand relative to the transmembrane domain (TMD) of the receptor differs substantially between the two templates PTH1R-LA-PTH and RAMP-bound CLR-CGRP (or AM1R complex) structures (Fig. S10B3). Both RAMP as well as different ligand conformations can cause these differences. The PTH1R and CLR are both similar in the receptor-ECD sequences and especially in their structural folding (Fig. S10A, S10B4), but the bound ligands (that are structurally known so far) are distinct in their secondary structures and overall arrangement to the receptor-ECD (with the ligand C-terminus) and to the transmembrane region (with the ligand N-terminus).

In the case of PTH1R, the LA-PTH ligand forms a straight helix from the ECD to the TMD at the receptor. In contrast, the bound CGRP or adrenomedullin in the CLR or AM1R, respectively, is kinked or bent in each case, and the C-terminus is unfolded compared to the straight LA-PTH helix (Fig. S10 B1-B2, B4). Notably, the ECD folding of the receptor is similar when comparing the structures of apo-ECD, RAMP1-CLR or CLR-RAMP1-CGRP complexes (Fig. S10B5), which is not indicative of global structural changes in the ECD folding of the receptor due to the binding of ligands or RAMPs.

Due to the specific orientation of the CLR-ECD and the differences in the length of extracellular loop 2 (EL2), the N-terminus of the CGRP ligand is different compared to LA-PTH at the transmembrane receptor part (Fig. S10B3 upper panel). Thus, the CGRP ligand is shifted towards TM1 and TM2 of the CLR compared to the LA-PTH orientation in the PTH1R.

Fourth, analysis of amino acid contacts between RAMP2 and the CLR ECD without ligand (Fig. S11B left) shows that essential amino acids in the receptor ECD that contact RAMP2 are also present in the N-terminal extracellular PTH1R sequence (Fig. S11A), e.g. amino acids Q45 and E54 (similar to Q54 in CLR). This fact supports the possibility that the RAMP2 ECD also binds to the PTH1R ECD in a mode comparable to that observed for the CLR-ECD-RAMP1 complex. In addition, corresponding and identical receptor amino acids in the ECD, such as R162, are involved in the binding of LA-PTH to PTH1R and CGRP to CLR, respectively (Fig. S10A and B middle-right parts), which also suggests a certain level of conservation in the ligand binding mode despite the differences in ligand conformation.

Finally, in the CLR-CGRP-RAMP1 complex, a distinct separation between receptor-ECD parts involved in either ligand or RAMP binding can be distinguished (Fig. S10B2), with corresponding amino acids involved

in RAMP1 binding at the CLR also found in the PTH1R-ECD.
Therefore, binding of the RAMP2-ECD to the PTH1R-ECD in a manner similar to that observed for the CLR-RAMP1-CGRP complex should in principle be possible.
Based on these observations, RAMP2 was structurally mapped to the structure of the PTH1R/ligand/G protein complex in two different ways.
On one hand, the RAMP2-ECD was bound to the PTH1R-ECD similarly to the CLR-CGRP-RAMP1 complex, retaining the entire PTH1R/LA-PTH structure already determined (Fig. S12A). For this aim, the superimposed (similar) receptor ECDs were used. The CLR-ECD apo-RAMP2 complex was superimposed on the PTH1R complex and the RAMP2-ECD was inserted into the PTH1R structure. The RAMP1-helix and the adjacent extracellular linker (Fig. S10A, B2) were additionally added into the PTH1R complex structure after superimposition of both complexes (Fig. S10B3). The RAMP1 sequence was substituted by the RAMP2 sequence (Fig. S10A) and the linker was manually connected to the RAMP2 ECD already merged into the initial PTH1R-ligand-RAMP2 model (Fig. S12A). Further the LA-PTH sequence was substituted by the PTH or by the PTHrP sequences, resulting in two initial homology models of the PTH1R-RAMP2-Gs complex with both ligands. These rough homology models were generated with the software SYBYL-X 2.0 (Certara, NJ, US) and optimized by energy minimization with the Amber99 force field until converging at a termination gradient of 0.05 kcal/mol* under constrained backbone atoms, followed by a 2 ns molecular dynamics simulation (MD) of side chains with constraint backbone atoms, with the exception of an un-constraint RAMP2 linker region. The entire complex was then energetically minimized without any constraint.
The second complex was generated by using the CLR-CGRP-RAMP1 complex[51] (PDB ID: 6e3y) as a template for arrangements between RAMP2 toward PTH1R, and the receptor ECD toward the receptor TMD. The PTH1R ECD bound with the LA-PTH C-terminus (amino acids 22-34) was separated from the entire complex[16] (PDB ID: 6nbh) and oriented towards the TM domain as supposed by the CLR-CGRP-RAMP1 complex. The entire RAMP1 was additionally inserted from this complex into the PTH1R model, and the sequence was substituted with the RAMP2 sequence. Finally, the two ligand fragments were manually connected between the extracellular C-terminus and the bound fragment (A1-R21) in the TM region. The ligand sequence was then substituted either by the PTH or the PTHrP sequences, respectively (Fig. S11A). These two models were optimized by energy minimization with the Amber99 force field until converging at a termination gradient of 0.05 kcal/mol* under constrained backbone atoms, followed by a 2 ns MD with constraint backbone atoms, except for the ligand region between AA S17-V21 (Fig. S11A). Again, the entire complexes were energetically minimized without any constraint.

#### Supplementary figures

**A**

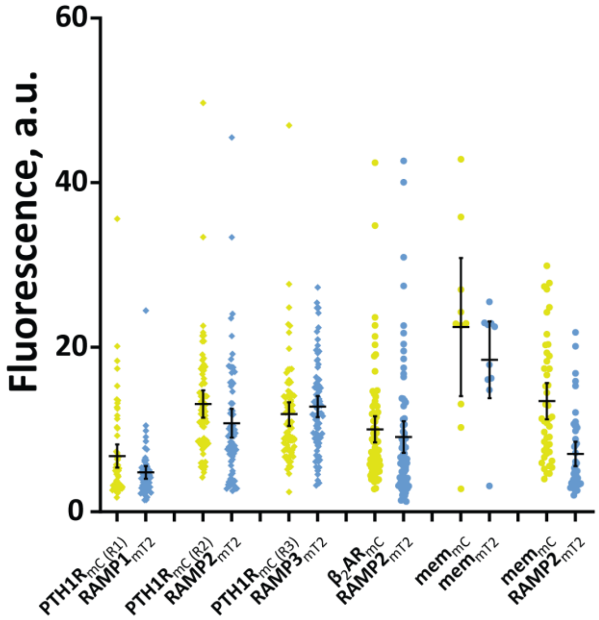


**Fig. S1.**

**Comparison of expression levels of tagged constructs used in intermolecular FRET photobleaching experiments.**

**(A)** Basal fluorescence emissions of mCitrine (mC, acceptor, yellow) and mTurquoise2 (mT2, donor, cyan) before photobleaching; the experimental setting corresponds to Fig. 1A. The data show individual values, mean and SD of at least three independent experiments.

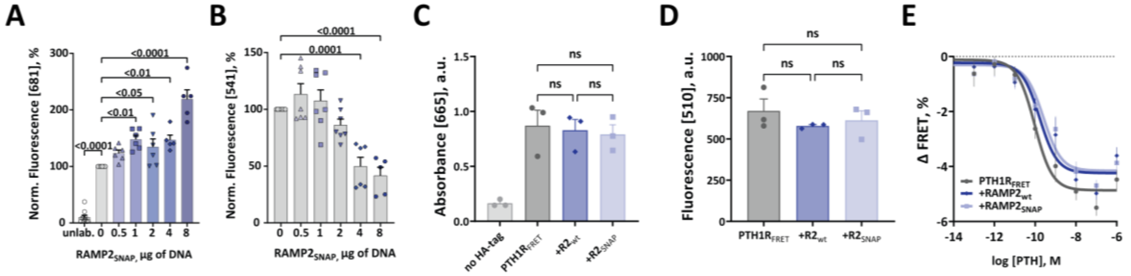


**Fig. S2.**

**Surface and total expression of PTH1R$_{FRET}$ biosensor are not modulated by RAMP2$_{SNAP}$ overexpression.**

**(A - B)** HEK293 cells were transfected with the PTH1R$_{FRET}$ biosensor plus different amounts of cDNA coding for RAMP2$_{SNAP}$. Emissions of SNAP-tag labelled with the 1 µM SNAP-Cell SiR-647 **(A)** and mCitrine **(B)** were collected in a plate reader. Bars represent means ± SEM, points are means of the duplicates of individual wells from three **(A)** and five **(B)** independent experiments. Significance between the groups was tested with one-way ANOVA followed by Dunnett's multiple comparisons test; ns: $p > 0.05$.

**(C - D)** HEK293 cells transiently expressing the PTH1R$_{FRET}$ biosensor were co-transfected with an empty control vector, RAMP2$_{wt}$ or RAMP2$_{SNAP.}$ **(C)** Comparison of cell surface expression levels of PTH1R$_{FRET}$ visualized by detecting the anti-HA tag epitope fused to its N-terminus and quantified by ELISA (absorbance at 665 nm). **(D)** Comparison of total expression levels of PTH1R$_{FRET}$ visualized by recording fluorescence of mCitrine in the same cells as in panel **(A)** The bars show means ± SEM of three independent experiments done in quadruplicates.

**(E)** Concentration-response curves obtained in HEK293 cells stably expressing the PTH1R$_{FRET}$ biosensor and co-transfected with RAMP2$_{wt}$ or RAMP2$_{SNAP.}$ Cells were stimulated with increasing concentrations of PTH. Curve fitting gave pEC$_{50}$ values for: PTH1R$_{FRET}$ = 10.02 ± 0.13, +RAMP2$_{wt}$ = 9.78 ± 0.19, and +RAMP2$_{SNAP}$ = 9.59 ± 0.20. The data show means ± SEM from three independent experiments done in quadruplicates.

Significance between the groups in panels **(D, E)** was tested by one-way ANOVA, followed by Tukey's multiple comparisons test; ns: $p > 0.05$.

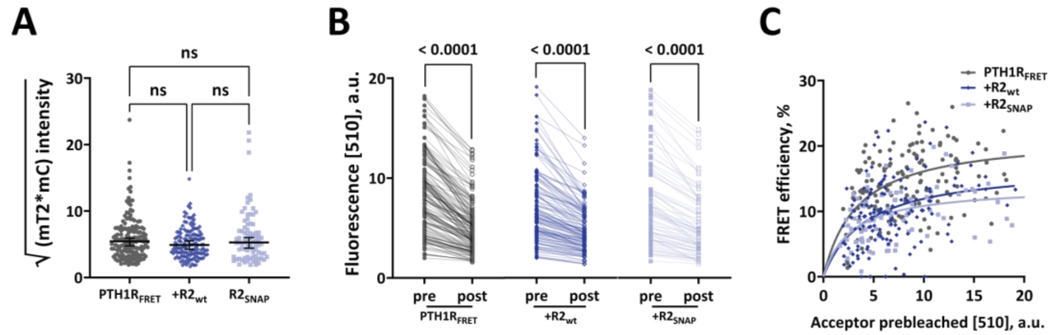


**Fig. S3.**

**Comparison of expression levels of fluorophores and photobleaching experiments with PTH1R$_{FRET}$ in intact HEK293 cells.**

**(A)** Basal fluorescence emission of mCitrine (mC, acceptor) and mTurquoise2 (mT2, donor) were measured in a confocal microscope before photobleaching of HEK293 cells expressing the indicated constructs. The square root of the product of mT2 and mC normalizes for different expression levels of fluorophores in order to compare biosensor expression between experimental groups. The data show median emission + 95 % CI from all cells examined from four independent experiments. Each data point represents a single cell. Significance between experimental groups was determined by Kruskal-Wallis nonparametric test with Dunn´s post-hoc test.

**(B)** Fluorescence emissions before (pre) and after (post) photobleaching shows comparable extents of photobleaching in the different experimental groups. Median photobleaching was PTH1R$_{FRET}$ = 39.9 %, RAMP2$_{wt}$ = 31.7 % and RAMP2$_{SNAP}$ = 28.5 %. Significance between pre and post emission was tested with Wilcoxon paired test, ns > 0.05.

**(C)** FRET efficiencies of PTH1R$_{FRET}$ in the absence or presence of RAMP2$_{wt}$ or RAMP2$_{SNAP}$ were calculated as described in Fig. 1. The data are plotted as a function of the emission of the acceptor before photobleaching. The curves were fitted with a one site-specific binding fit. Each data point represents a single cell. Data are from the following numbers of cells obtained in four independent experiments: PTH1R$_{FRET}$ (n = 120), +RAMP2$_{wt}$ (n = 96), +RAMP2$_{SNAP}$ (n = 72).

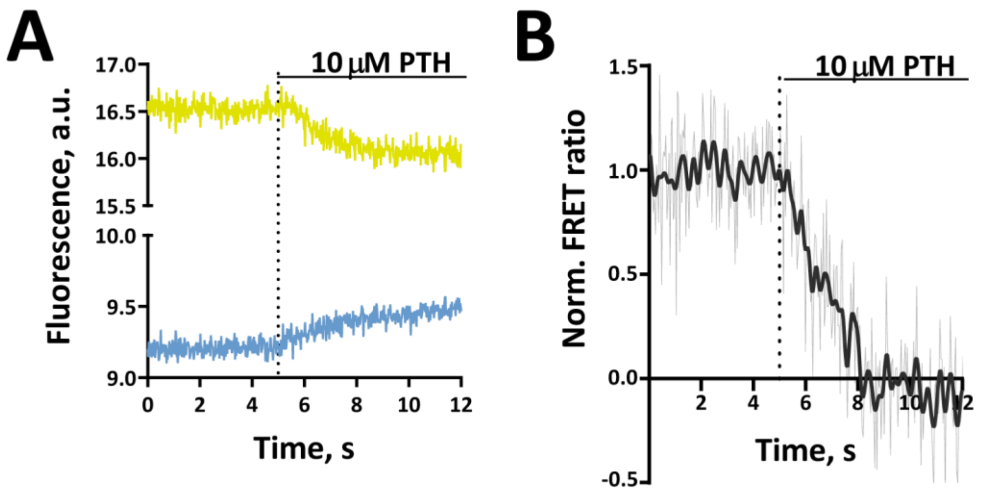


**Fig. S4.**

**PTH-induced activation changes recorded with the $PTH1R_{FRET}$ biosensor.**

Representative FRET traces of PTH-mediated changes in intramolecular FRET in single HEK293 cells stably expressing of $PTH1R_{FRET}$. Horizontal lines indicate application of 10 μM PTH with a rapid superfusion system. **(A)** Traces of donor (mT2) and acceptor (mC) fluorescence. **(B)** FRET ratio calculated from **(A)**. Traces were normalized to the baseline (set to 1) and plateau after stimulation (set to 0). Shown are FRET ratio traces raw (grey) and Fourier-lowpassed (black). Traces are representative of n = 41 cells, acquired in five independent experiments.

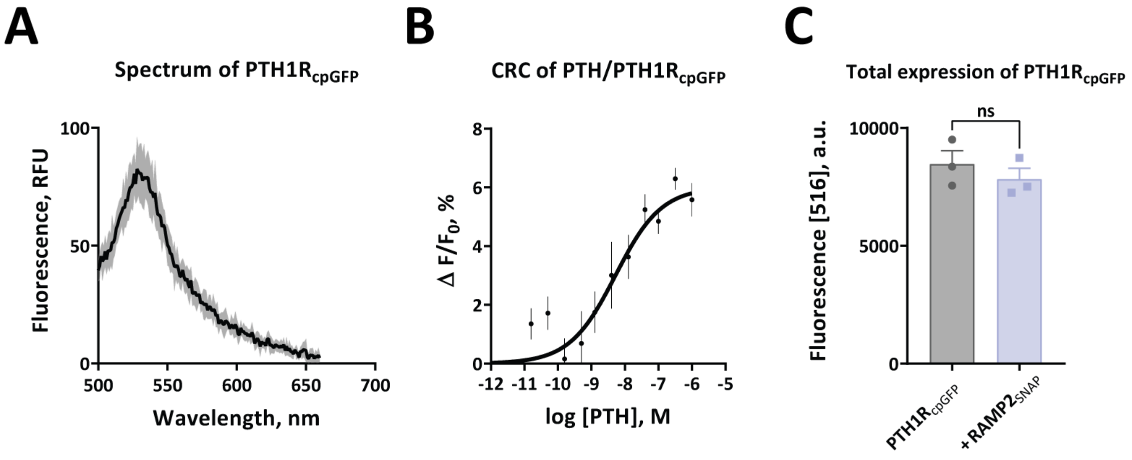


**Fig. S5.**

**Spectral and pharmacological characterization of PTH1R$_{cpGFP}$.**

Plate reader experiments with HEK293 cells transiently expressing the PTH1R$_{cpGFP}$ biosensor.

**(A)** Fluorescence emission spectra of PTH1R$_{cpGFP}$ upon excitation at 460 nm in the presence of 10 µM PTH. The data are from two independent experiments done in quadruplicate and show mean ± SEM.

**(B)** Concentration-response curve for stimulation with increasing concentrations of PTH. The data show mean ± SEM of three independent experiments performed in quadruplicate. The curve was fitted with a three-parameter concentration-response curve fit and gave a pEC$_{50}$ ± SEM of 8.28 ± 0.27.

**(C)** Comparison of total expression levels for PTH1R$_{cpGFP}$ in the absence and presence of RAMP2$_{SNAP}$. Shown are means ± SEM of three independent experiments, where each individual mean was calculated from 48 wells from a single 96-well plate. A t-test was used to assess significance between the groups; ns > 0.05.

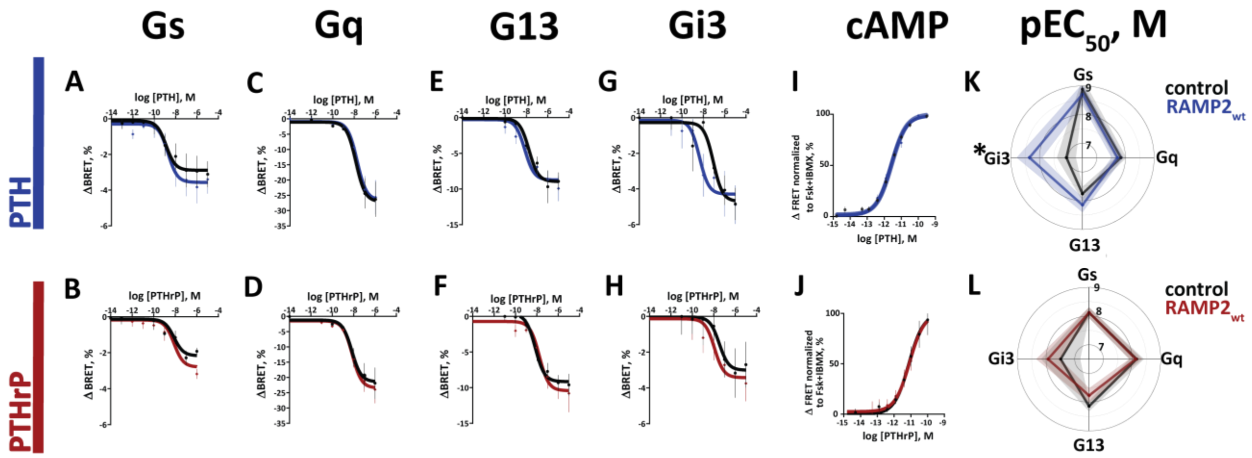


**Fig. S6.**

**RAMP2 effects on PTH-stimulated G protein activation.**

(**A - H**) HEK293 cells transiently transfected with cDNA encoding for BRET biosensors of: $G_s$ (**A, B**), $G_q$ (**C, D**), $G_{13}$ (**E, F**), $G_{i3}$ (**G, H**) along with $PTH1R_{wt}$, with or without $RAMP2_{wt}$.

(**I, L**) HEK293 cells transiently transfected with cDNA encoding for the cAMP-based FRET biosensor (Epac-$S^{H187}$), along with $PTH1R_{wt}$, with or without $RAMP2_{wt}$.

BRET signals were recorded in a plate reader from cells stimulated with PTH (black, blue) or PTHrP (black, red). Shown are time courses of agonist stimulation and corresponding concentration-response curves, fitted with a three-parameter concentration-response curve fit. Data are means ± SEM of at least n = 3 independent experiments performed in duplicates or more. For further statistics and results see SI Appendix, Table S1 and S2.

(**K, L**) "Spider plots" showing mean ± SEM $pEC_{50}$ (M) values calculated from the concentration-response curves.

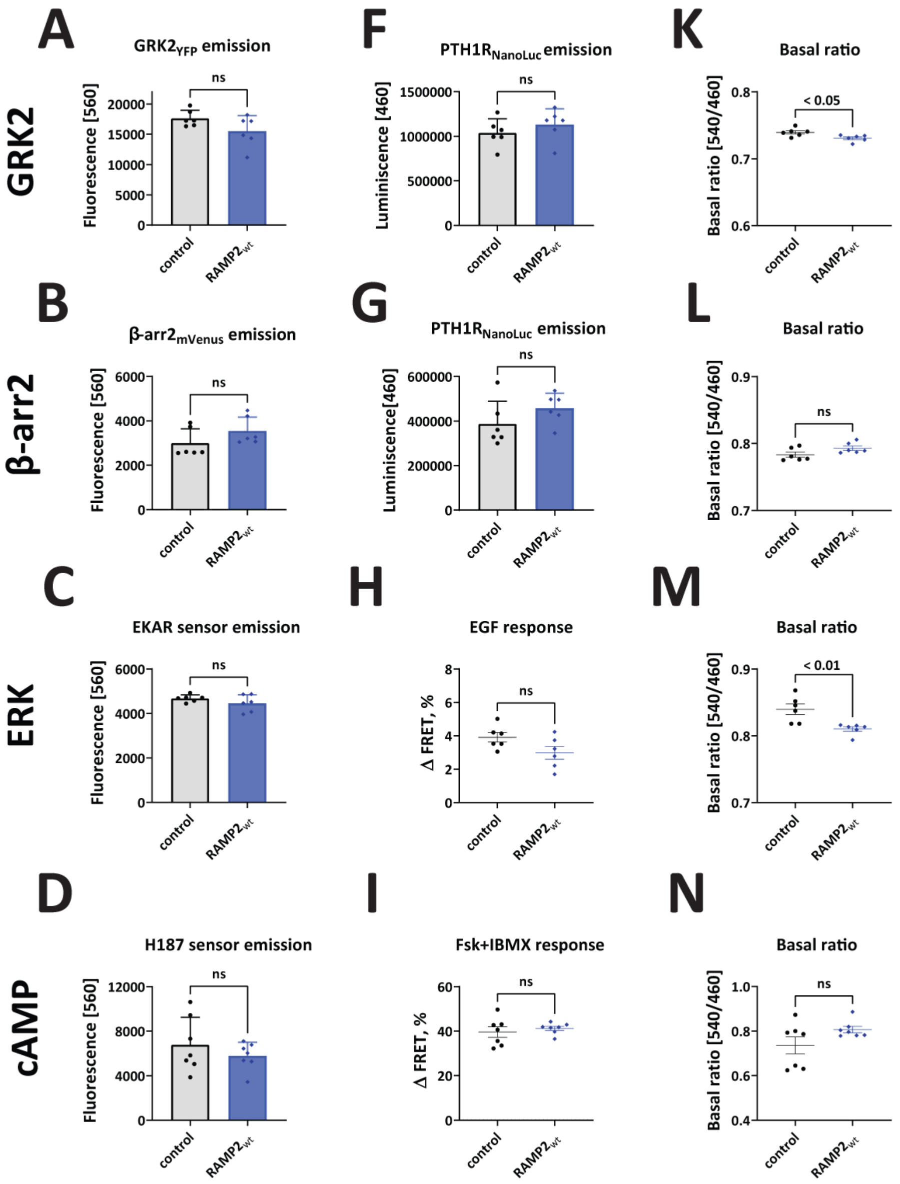


**Fig. S7.**

**Expression controls for non-G protein functional assays.**

Comparison of expression levels of constructs used in non-G protein PTH1R functional assays in the absence (grey) and presence (blue) of $RAMP2_{wt}$. Data were obtained in plate reader experiments with HEK293 cells transiently expressing a combination of the indicated constructs. Fluorescence emissions indicate: at 560 nm the expression level of **(A)** $GRK2_{YFP}$, **(B)** β-arrestin$2_{mVenus}$, **(C)** EKAR sensor and **(D)** cAMP biosensor H187, at 460 nm the expression level of **(F - G)** $PTH1R_{NanoLuc}$. **(H)** Comparison of ERK responses to 100 ng/mL Epidermal growth factor (EGF). **(I)** cAMP responses to 10 µM forskolin plus 100 µM IBMX. **(K - N)** Comparison of basal ratios before stimulation of each construct in presence of absence of $RAMP2_{wt}$. Shown are means ± SEM of at least three independent experiments. A t-test was used to assess significance between the groups, ns > 0.05.

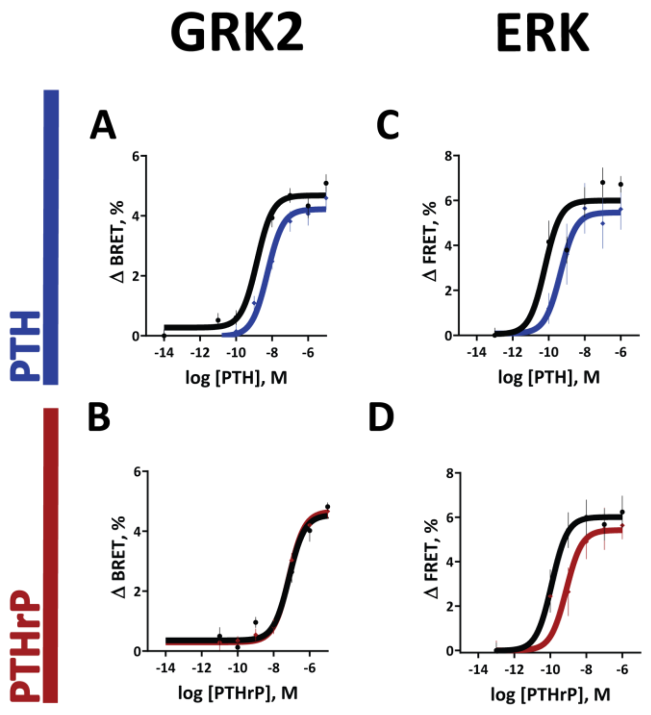


**Fig. S8.**

**RAMP2 effects on non-G protein signalling**.

**(A - D)** HEK293 cells were transiently transfected with cDNA encoding for: $GRK2_{YFP}$ (A, B) along with $PTH1R_{NanoLuc}$, with or without $RAMP2_{wt}$, or with EKAR biosensor (C - D) along with $PTH1R_{wt,}$ with or without $RAMP2_{wt}$. BRET signals were recorded in a plate reader from cells stimulated with PTH (black, blue) or PTHrP (black, red). Shown are time courses of agonist stimulation and corresponding concentration-response curves, fitted with a three-parameter concentration-response curve fit. Data are means ± SEM of at least n = 3 independent experiments performed in quadruplicates or more. For further statistics and results see SI Appendix, Table S3 and S4.

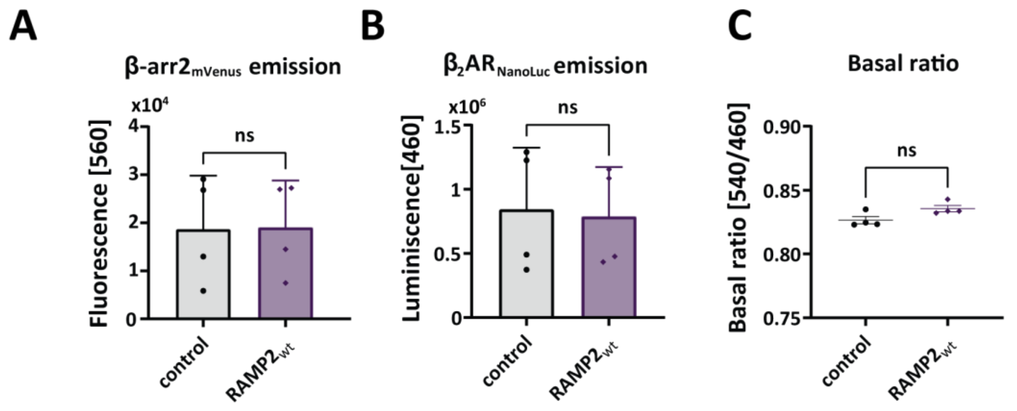


**Fig. S9.**

**Expression controls for β-arrestin2 recruitment to $\beta_2$AR.**

**(A, B)** Comparison of expression levels of constructs used in the experiments for isoprenaline-induced β-arrestin2 recruitment to $\beta_2$AR.

**(C)** Comparison of basal ratios before stimulation. All data are from four independent experiments done in quadruplicates and represent means ± SEM where each individual mean was calculated from 48 wells in a single 96-well plate. A t-test was used to assess significance between the groups, ns > 0.05.

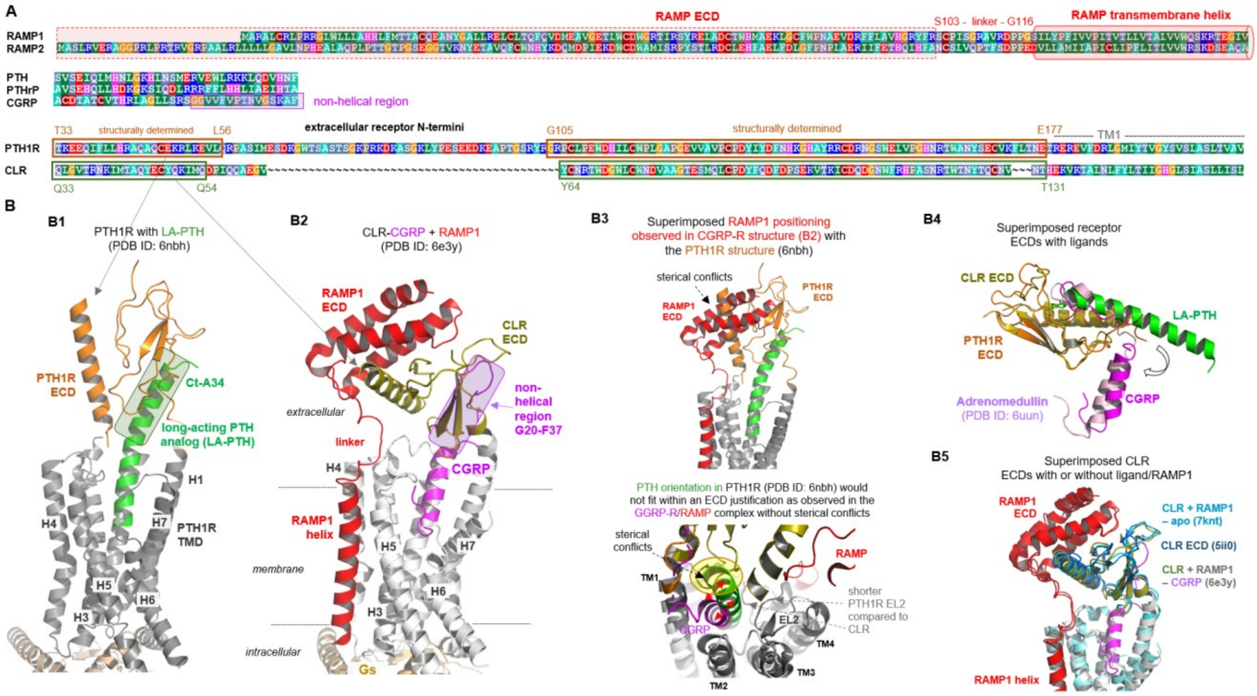


**Fig. S10.**

**Sequence and structural information used to construct PTH1R ligand-RAMP2 complexes - Part I.**

**(A)** Sequence comparison between RAMP1 and RAMP2, ligands PTH, PTHrP, CGRP, and receptor ECDs, with highlighted dimensions in length or structural features. The alignment was visualized using the software BioEdit[67]. Specific background colors reflecting chemical properties of the amino acid side chains or the type of amino acid: black-proline; blue-positively charged; cyan/green-aromatic and hydrophobic; green- hydrophobic; red-negatively charged; gray-hydrophilic; dark red-cysteines; and magenta-histidine.

**(B)** Structural templates for model building are visualized, including the PTH1R complex structure (B1) and the CLR-CGRP-RAMP1 complex (B2). The PDB ID's are annotated above the structures. In B3 both complexes are superimposed to highlight differences and similarities. In addition, determined receptor ECD-ligand complexes are superimposed in B4 to visualize conservation of the ECD folds, but also to show differences in the bound ligand conformations. Of note, the receptor ECDs with or without ligand and RAMP (B5) share a similar global fold, which indicates a low impact of ligand or RAMP binding on the receptor ECD core structure.

H: helix
EL extracellular loop
IL: intracellular loop
ECD: extracellular domain

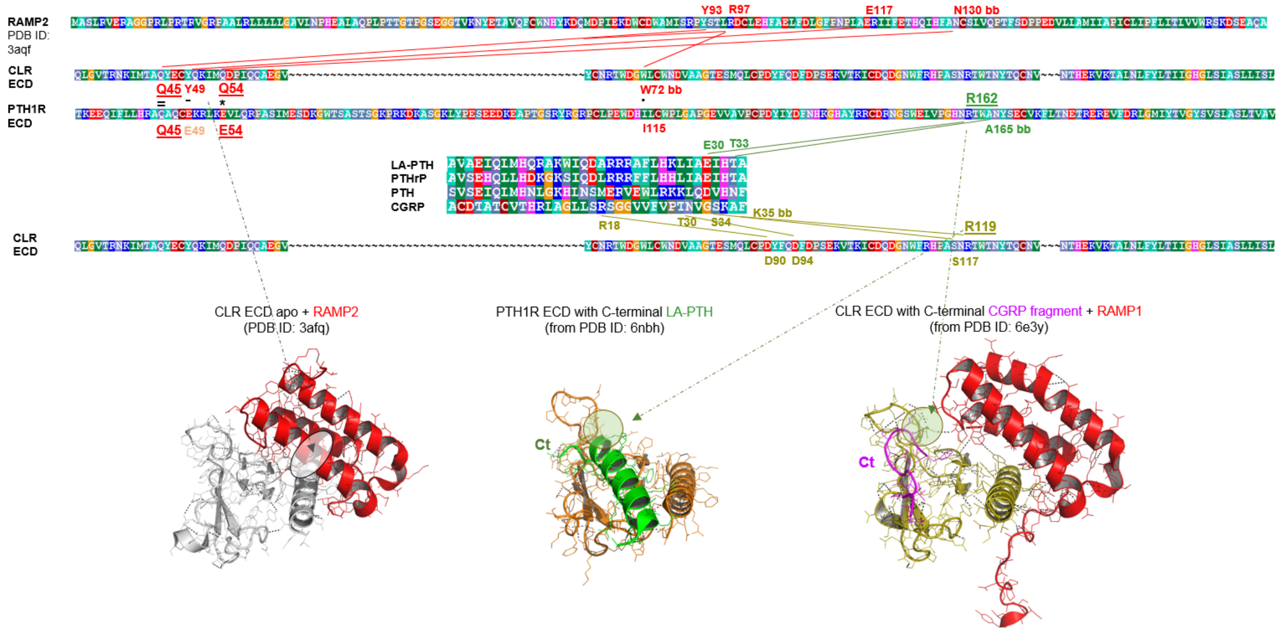


**Fig. S11.**

**Sequence and structural information used to construct PTH1R ligand-RAMP2 complexes - Part II.**

Analyses of interactions (hydrogen-bond contact distances) between RAMP2 and the CLR ECD (red lines indicating feasible interactions) reveal few essential hydrophilic contacts and similar amino acids of the CLR ECD can also be found at corresponding positions in the sequence of the PTH1R ECD (e.g. Q45 in the central contact region between receptor ECD and RAMP ECD). Moreover, ligand contacts between the ECD's of both receptors are shared by corresponding amino acids as R162, even the ligand conformation (C-terminus) is different in the bound states.

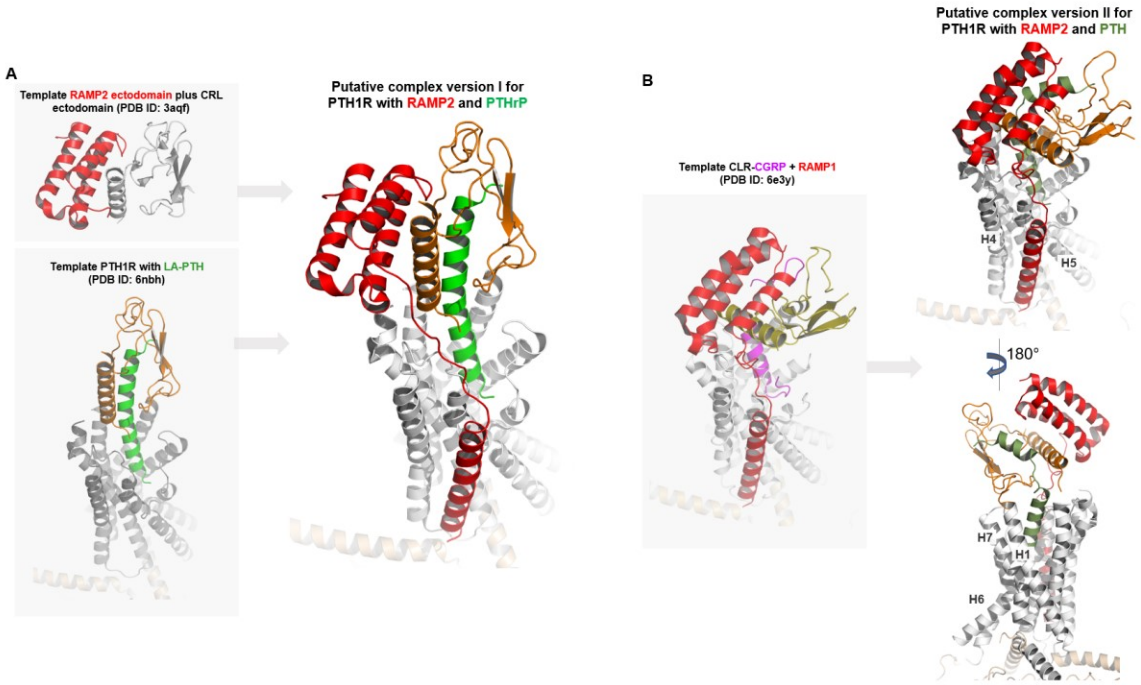


**Fig. S12.**

**Putative structural PTH1R-ligand-RAMP2-Gs complexes.**

Starting with appropriate structural templates to model putative PTH1R-RAMP-ligand complexes (PDB ID's: 6nbh, 3aqf)[16,51,68] resulted in two versions of feasible arrangements. **(A)** In version I the RAMP2 ECD is oriented toward the PTH1R, but the principle receptor ECD is adjusted according to the already determined PTH1R-LA-PTH-Gs complex, whereby the ligand forms a regular straight helix. **(B)** Taking the already solved CLR-CGRP-RAMP1-Gs complex[51] as a structural template (PDB ID: 6e3y) to adjust the complex components relative to each other, the PTH1R ECD bound with RAMP2 ECD is differently oriented toward the TMD and the ligand (e.g. PTH) must have a modified secondary structure in the central part compared to LA-PTH in the determined PTH1R complex **(A)**.

#### Supplementary tables

**Table S1: Potency ($pEC_{50}$) and $E_{max}$ values for PTH-induced G protein activation.**

| | | PTH | | | |
|---|---|---|---|---|---|
| | | **control** | **RAMP2$_{wt}$** | ***p*** | ***n*** |
| **$G_q$ activation** | **$pEC_{50}$** | 7.84 ± 0.19 | 7.72 ± 0.20 | ns | 3 |
| | **$E_{max}$** | 26.02 ± 2.38 | 25.63 ± 2.35 | ns | |
| **$G_s$ activation** | **$pEC_{50}$** | 8.88 ± 0.39 | 8.71 ± 0.36 | ns | 4 |
| | **$E_{max}$** | 2.78 ± 0.43 | 3.28 ± 0.45 | ns | |
| **$G_{13}$ activation** | **$pEC_{50}$** | 7.76 ± 0.24 | 8.15 ± 0.25 | ns | 4 |
| | **$E_{max}$** | 8.84 ± 0.90 | 8.47 ± 0.90 | ns | |
| **$G_{i3}$ activation** | **$pEC_{50}$** | 7.06 ± 0.33 | 8.35 ± 0.44 | < 0.05 | 4 |
| | **$E_{max}$** | 4.43 ± 0.68 | 4.16 ± 0.73 | ns | |
| **cAMP accumulation** | **$pEC_{50}$** | 11.57 ± 0.05 | 11.60 ± 0.06 | ns | 4 |
| | **$E_{max}$** | 100.00 ± 2.46 | 99.18 ± 2.86 | ns | |

Potency ($pEC_{50}$, M) and $E_{max}$ (%) values were obtained from plate reader experiments as shown in Fig. S6. Data are means ± SEM of n independent experiments. Extra-sum-of-squares test was used to assess the difference between the curves, ns > 0.05.

**Table S2: Potency ($pEC_{50}$) and $E_{max}$ values for PTHrP-induced G protein activation.**

| | | PTHrP | | | |
|---|---|---|---|---|---|
| | | **control** | **$RAMP2_{wt}$** | ***p*** | ***n*** |
| **$G_q$ activation** | **$pEC_{50}$** | 8.16 ± 0.25 | 8.13 ± 0.22 | ns | 3 |
| | **$E_{max}$** | 20.20 ± 2.29 | 21.98 ± 2.21 | ns | |
| **$G_s$ activation** | **$pEC_{50}$** | 8.12 ± 0.18 | 8.12 ± 0.18 | ns | 4 |
| | **$E_{max}$** | 2.04 ± 0.27 | 2.62 ± 0.27 | < 0.05 | |
| **$G_{13}$ activation** | **$pEC_{50}$** | 8.14 ± 0.18 | 7.77 ± 0.25 | ns | 4 |
| | **$E_{max}$** | 9.48 ± 0.74 | 9.72 ± 1.01 | ns | |
| **$G_{i3}$ activation** | **$pEC_{50}$** | 7.49 ± 0.68 | 7.93 ± 0.41 | ns | 4 |
| | **$E_{max}$** | 2.98 ± 0.82 | 3.29 ± 0.58 | ns | |
| **cAMP accumulation** | **$pEC_{50}$** | 11.12 ± 0.14 | 11.08 ± 0.19 | ns | 4 |
| | **$E_{max}$** | 100.00 ± 8.60 | 97.40 ± 11.03 | ns | |

Potency ($pEC_{50}$, M) and $E_{max}$ (%) values were obtained from plate reader experiments as shown in Fig. S6. Data are means ± SEM of n independent experiments. Extra-sum-of-squares test was used to assess the difference between the curves, ns > 0.05.

**Table S3: Potency ($pEC_{50}$) and $E_{max}$ values for PTH-induced downstream effects.**

| | | PTH | | | |
|---|---|---|---|---|---|
| | | **control** | **RAMP2$_{wt}$** | ***p*** | ***n*** |
| **GRK2 recruitment** | **$pEC_{50}$** | 8.84 ± 0.13 | 8.25 ± 0.14 | < 0.05 | 3 |
| | **$E_{max}$** | 4.40 ± 0.23 | 4.23 ± 0.25 | ns | |
| **β-arrestin2 recruitment** | **$pEC_{50}$** | 9.45 ± 0.14 | 9.21 ± 0.12 | ns | 3 |
| | **$E_{max}$** | 3.66 ± 0.21 | 6.48 ± 0.32 | < 0.05 | |
| **ERK phosphorylation** | **$pEC_{50}$** | 10.21 ± 0.24 | 9.34 ± 0.39 | ns | 3 |
| | **$E_{max}$** | 5.95 ± 0.76 | 5.39 ± 1.00 | ns | |

Potency ($pEC_{50}$, M) and $E_{max}$ (%) values are obtained from plate reader experiments as shown in Fig. 5 and Fig. S8. Data are means ± SEM of n independent experiments. Extra-sum-of-squares test was used to assess the difference between the curves, ns > 0.05.

**Table S4: Potency ($pEC_{50}$) and $E_{max}$ values for PTHrP-induced downstream effects.**

| | | PTHrP | | | |
|---|---|---|---|---|---|
| | | **control** | **RAMP2$_{wt}$** | ***p*** | ***n*** |
| **GRK2 recruitment** | **$pEC_{50}$** | 7.04 ± 0.12 | 7.23 ± 0.11 | ns | 3 |
| | **$E_{max}$** | 4.25 ± 0.24 | 4.32 ± 0.20 | ns | |
| **β-arrestin2 recruitment** | **$pEC_{50}$** | 8.56 ± 0.12 | 8.45 ± 0.09 | ns | 3 |
| | **$E_{max}$** | 3.45 ± 0.16 | 5.85 ± 0.21 | < 0.05 | |
| **ERK phosphorylation** | **$pEC_{50}$** | 9.92 ± 0.40 | 9.12 ± 0.49 | ns | 3 |
| | **$E_{max}$** | 6.02 ± 0.82 | 5.42 ± 0.78 | ns | |

Potency ($pEC_{50}$, M) and $E_{max}$ (%) values were obtained from plate reader experiments as shown in Fig. 5 and Fig. S8. Data are means ± SEM of n independent experiments. Extra-sum-of-squares test was used to assess the difference between the curves, ns > 0.05.

**Table S5: Potency ($pEC_{50}$) and $E_{max}$ values for isoprenaline-induced β-arrestin2 recruitment.**

| | | Isoprenaline | | | |
|---|---|---|---|---|---|
| | | **control** | **$RAMP2_{wt}$** | ***p*** | ***n*** |
| **β-arrestin2 recruitment** | **$pEC_{50}$** | 7.58 ± 0.13 | 7.81 ± 0.15 | ns | 4 |
| | **$E_{max}$** | 2.43 ± 0.10 | 2.72 ± 0.13 | ns | |

Potency ($pEC_{50}$, M) and $E_{max}$ (%) values were obtained from plate reader experiments as shown in Fig. 5. Data are means ± SEM of n independent experiments. Extra-sum-of-squares test was used to assess the difference between the curves, ns > 0.05.
